# Environment-driven active transport of influenza A virus

**DOI:** 10.64898/2026.09.03.749308

**Authors:** Siddhansh Agarwal, Liya F. Oster, Boris Veytsman, Greg Huber, Daniel A. Fletcher

## Abstract

Biological media such as airway mucus and extracellular matrix are usually viewed as transport barriers that particles cross by passive diffusion or with internal engines. We show instead that a particle can move actively by modifying the landscape it traverses, creating environmental memory and directional cues in two and three dimensions. Influenza A virus (IAV) realizes this principle through its envelope proteins hemagglutinin (HA) and neuraminidase (NA), which bind and cleave sialylated glycan receptors, respectively. Combining theory, simulations and single-virus tracking, we connect bind–cleave kinetics and HA–NA organization to macroscopic transport. Cleavage dissipates chemical free energy, biases rebinding to the edited landscape and leaves a trail that shapes future encounters. In heterogeneous receptor landscapes, multivalent binding biases motion toward higher receptor density, while receptor destruction by NA can amplify this bias by sharpening the contrast sampled by HA. Experiments on reconstituted glycan membranes show that IAV steps are biased up local receptor gradients, as predicted. The theory suggests that virion-to-virion variability can distribute transport functions across a population, providing a physical hedge against complex receptor environments. Together, these results establish environment-driven active matter as a mechanism for motorless transport powered and guided by chemical modification of the environment.

---

How can a nanoscale particle with no apparent motor move directionally through a sticky, crowded environment such as the mucus layer in the lungs? Small particles are usually considered passive diffusers or active particles driven by catalytic surfaces, phoretic flows or response circuits [1–4]. Diffusive exploration can be reshaped by prescribed spatial disorder or by rules that depend on past motion [5–7]. A landscape that the particle modifies as it moves provides a third route by coupling reversible attachment to an irreversible change that dissipates chemical free energy and biases subsequent binding [8–10]. The environment becomes part of the mechanism of motion, defining a class of *environment-driven active matter*.

Influenza A virus (IAV) provides a realization of this idea. Before entering an airway epithelial cell, a virion must traverse receptor-bearing mucus and periciliary glycoproteins to reach the cell surface [11, 12]. Its two major envelope proteins form a minimal “bind–cleave” pair: hemagglutinin (HA) reversibly binds sialylated glycan receptors, whereas neuraminidase (NA) irreversibly cleaves them [13–15]. Two axes define this design space, spanning uniform to heterogeneous receptor distributions in the environment and mixed to polarized binder–cleaver organization on the particle (Fig. 1).

**Fig. 1.**
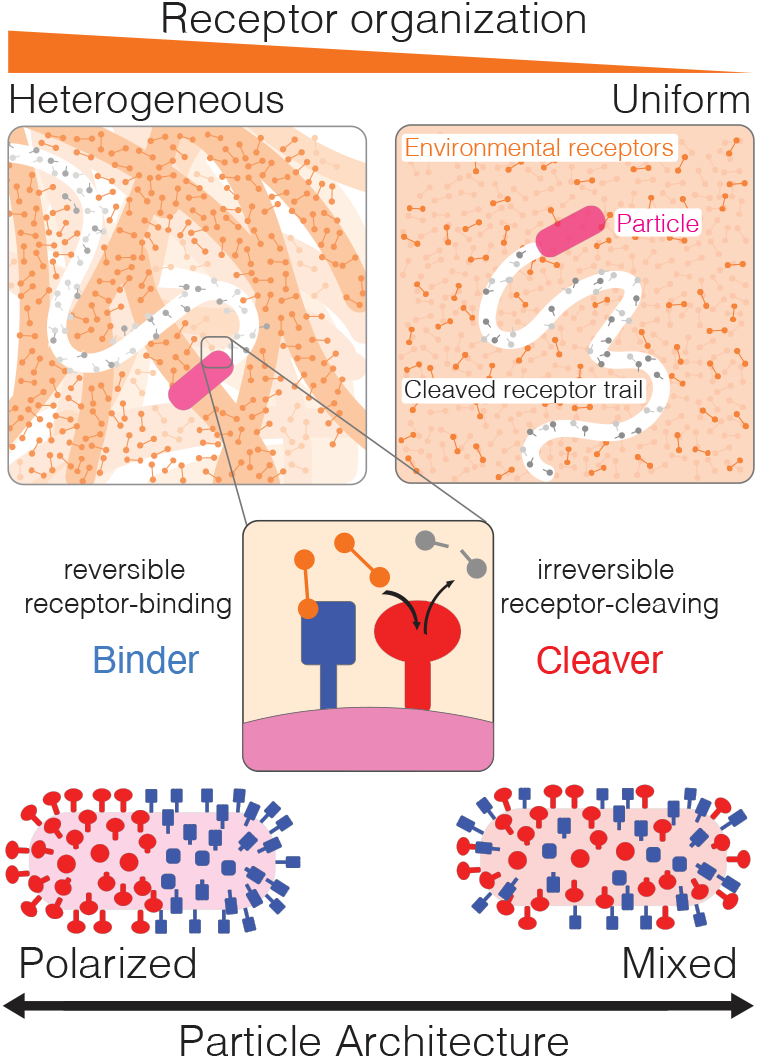
Receptor organization and particle architecture in environment-driven active matter. Receptor landscapes range from uniform to heterogeneous, while binders and cleavers are mixed or polarized on the particle. Reversible binding couples the particle to receptors; irreversible cleavage dissipates chemical free energy and records its path.

These axes are coupled because particle architecture determines where receptors are edited, and environmental organization sets what subsequent binding can read. Previous work showed that passive diffusion may be too slow for IAV to cross the mucus barrier before mucociliary clearance and that HA–NA chemistry can drive burnt-bridge-like locomotion by suppressing rebinding to used receptors [13, 14]. Related theories have described specific local modes of receptor-coupled surface motion [16, 17]. We ask how bind–cleave dynamics generate macroscopic transport in complex environments, where the outcome depends on whether the particle re-encounters its edits and on the receptor landscape’s pre-existing spatial structure. In 3D, a depleted region can be left behind; on a 2D surface, repeated returns make the written trail a memory that first extends exploration and ultimately promotes detachment. Across the airway, receptor density, affinity and accessibility vary, so the same machinery must preserve motion through sparse glycans, escape adhesive decoys and respond to receptor-rich cell surfaces.

These demands need not be met by one optimal particle. IAV particles vary significantly in morphology and HA–NA composition [18], and the nonlinear coupling between binding, cleavage and geometry can support transport states not represented by the mean phenotype. We combine stochastic simulations, mean-field theory and single-virus tracking experiments to connect binding, cleavage and particle architecture to transport, gradient guidance and complementary population functions.

## Multivalent attachment and cleavage control 3D diffusion

We begin with a uniform receptor environment, where any directional memory must be written by cleavage and read through subsequent binding (Fig. 1). To isolate this feedback, we extend our effective 1D model [14] to a rod that translates and rotates in 3D, coarse-graining transverse surface structure into *N*_tot_ = *N*_*b*_ + *N*_*c*_ axial binder and cleaver sites. Particle morphology, dimensions and total ligand number remain fixed, and we vary only the binder:cleaver ratio, ligand organization and the binding and cleavage parameters *K*_*D*_ and *K*_*C*_. Smaller *K*_*D*_ gives longer-lived attachments, whereas larger *K*_*C*_ accelerates receptor removal. Polarized particles segregate binders and cleavers, while mixed particles intersperse them (Fig. 2a; SI Section 1).

**Fig. 2.**
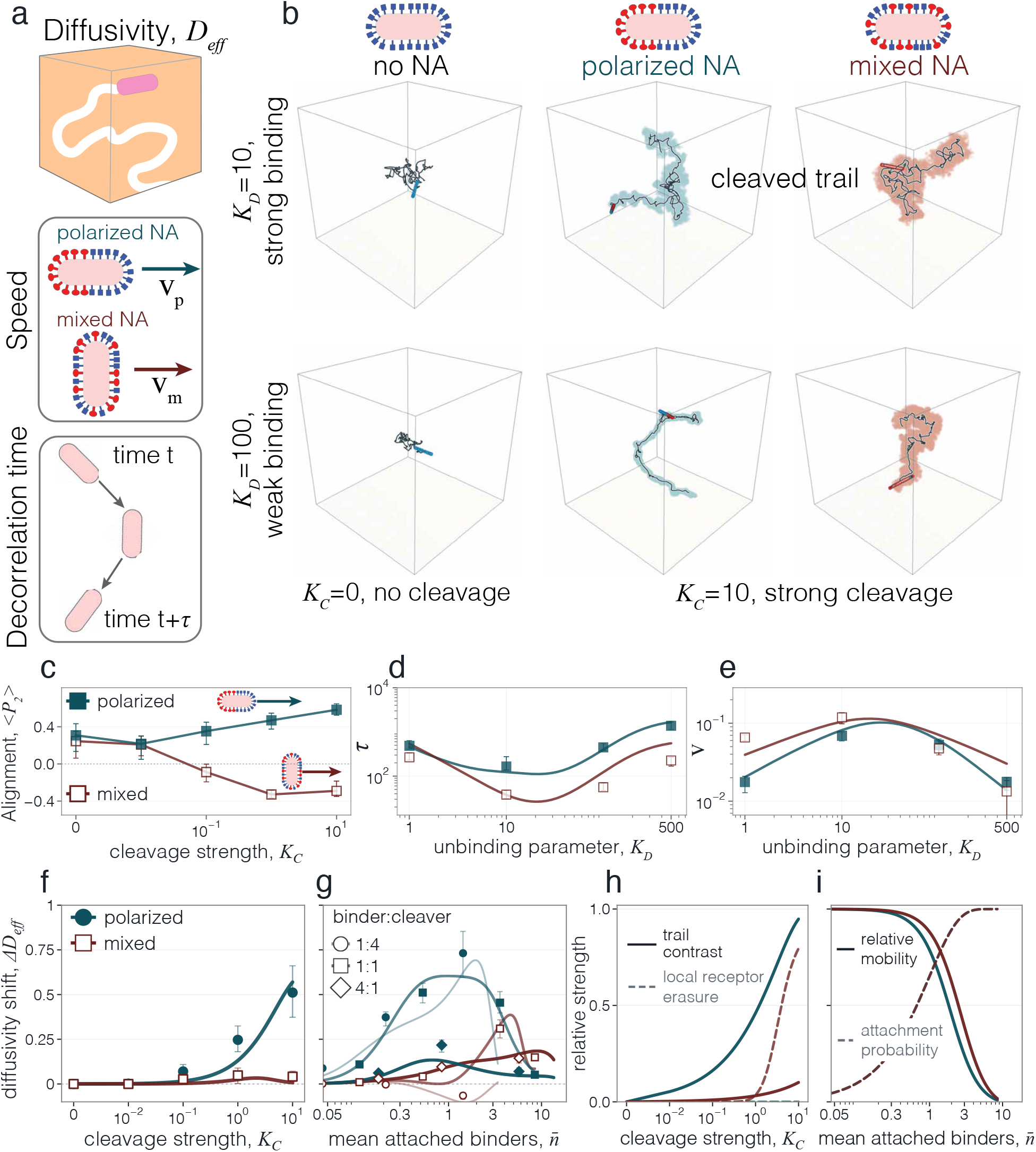
Multivalent attachment and local receptor depletion govern 3D exploration. **a**, Polarized (blue) and mixed (red) architectures in a uniform 3D receptor region. **b**, Trajectories at *K*_*C*_ = 0 and 10 for *K*_*D*_ = 10 and 100. Teal and red trails mark receptors cleaved by polarized and mixed particles, respectively. **c**, Body-axis alignment ⟨*P*_2_⟩ versus *K*_*C*_ ; positive and negative values indicate axial and transverse motion. **d**,**e**, Persistence time *τ* and receptor-driven speed *v* versus *K*_*D*_, with stronger binding to the left. **f**, Diffusivity shift, Δ*D*_eff_ = *D*_eff_ (*K*_*C*_ ) − *D*_eff_ (0), versus *K*_*C*_ . **g**, Δ*D*_eff_ versus the average number of attached binders 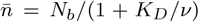, a measure of collective attachment, for binder:cleaver ratios 1:4, 1:1, and 4:1. **h**, Difference in remaining receptor density (solid) and probability of entering a locally depleted state (dashed); the latter vanishes for polarized particles. **i**, Fraction of free-particle mobility remaining after attachment (solid) and probability of at least one attachment (dashed). Symbols show simulations, lines in **c** guide the eye, and curves in **d–i** show mean-field predictions. Error bars are defined in Supplementary Information (SI) Section 2.

Without cleavage, repeated binding and unbinding produce compact trajectories at *K*_*D*_ = 10, whereas *K*_*D*_ = 100 approaches passive diffusion (Fig. 2b). At *K*_*C*_ = 10, polarized particles leave narrow teal trails of cleaved receptors and explore farther, especially at weaker binding. Mixed particles clear broader red footprints, with the *K*_*D*_ = 10 particle remaining mobile while local receptor depletion arrests the *K*_*D*_ = 100 particle within the region it has cleared. Cleavage can therefore extend transport or remove the intact receptors needed to sustain it. The body-axis alignment 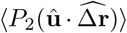 compares the displacement direction 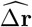 with the particle axis **û**, where *P*_2_ is the second Legendre polynomial. It becomes positive for polarized particles and negative for mixed particles, indicating axial and transverse motion, respectively, as *K*_*C*_ increases (Fig. 2c).

The distance covered during a directed run is set by *vτ* : cleavage generates the speed *v*, while rotational dynamics set the time *τ* over which that direction persists (Fig. 2d,e). Writing *a* = *p, m* for polarized and mixed architectures, respectively, each binder samples an effective number *ν* of receptors within range *α*, and attached binders resist rotation according to their lifetime *τ*_*b*_ and centered axial spread *I*_*b,a*_, giving

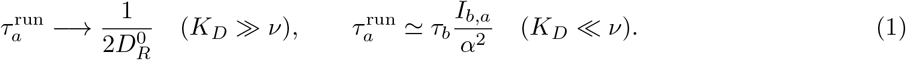

Weak binding recovers the free rotational-diffusion lifetime 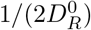, while strong binding makes persistence depend on attachment lifetime and binder placement. Repeated binding and unbinding produce the largest angular fluctuations between these limits and therefore the minimum in *τ* . Polarized particles retain a head– tail direction, whereas mixed particles are invariant under axis reversal, giving 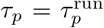 and 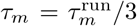 (SI Section 2).

Attachments also transmit force, and the axial or transverse direction identified by ⟨*P*_2_ ⟩determines which friction coefficient enters our earlier 1D speed balance [14]:

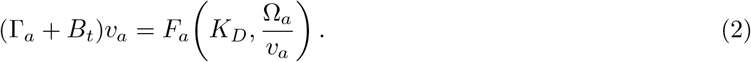

Here Γ_*a*_ is the corresponding background friction, and *B*_*t*_ is attachment-induced friction. The force *F*_*a*_ arises from the intact-receptor difference across the particle, while Ω_*a*_*/v*_*a*_ measures receptor removal per passage. Faster motion leaves less time for cleavage, so this balance recovers the previously derived 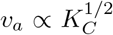 limit at weak cleavage [14]. Polarized segregation on the particle preserves intact receptors ahead of the binders, allowing *v*_*p*_ to rise with *K*_*C*_ before saturating. In mixed particles, the receptor difference first grows and then collapses as both sampled regions are depleted, making *v*_*m*_ non-monotonic. At fixed *K*_*C*_, weak attachment and strong-binding friction produce an intermediate-affinity maximum.

While the particle remains coupled to intact receptors, its speed and directional persistence contribute 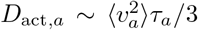 to long-time diffusion (SI Section 2). Built-in front–back asymmetry biases a polarized particle to move binder-first into intact receptors, whereas a mixed particle has no preferred direction and can reverse across the body-wide footprint it just depleted, losing all attachments until transverse motion brings intact receptors back within reach. Because a 3D walk need not return to older positions [19], this recent footprint dominates the loss of mobility. For architecture *a* = *p, m, f*_mob,*a*_ is the fraction of time the particle remains mobile and bound to receptors, and *D*_eff,*a*_(*K*_*C*_) is its long-time diffusivity at cleavage strength *K*_*C*_. The corresponding diffusivity without cleavage is *D*_bu,*a*_ ≡ *D*_eff,*a*_(0), so the diffusivity shift is

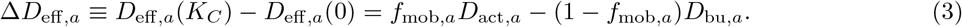

Spatial segregation of NA leaves intact receptors available to polarized particles, keeping *f*_mob,*p*_ ≃ 1. Their Δ*D*_eff,*p*_ remains positive and plateaus as *K*_*C*_ rises (Fig. 2f). For mixed particles, Δ*D*_eff,*m*_ peaks at intermediate *K*_*C*_ because strong cleavage depletes receptors along the particle and can eventually make the shift negative. Figure 2h shows this trade-off directly, with the receptor contrast crossing the locally depleted fraction 1 − *f*_mob,*m*_.

When binder number varies, the average number of attached binders, 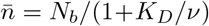, combines binder number and single-binder affinity into avidity (Fig. 2g). Too few attachments fail to transmit the receptor difference, while too many increase friction and suppress mobility (Fig. 2i). More binders can achieve the same 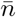 through weaker, shorter-lived attachments and therefore less attachment-induced friction. Decreasing the binder:cleaver ratio strengthens receptor removal but leaves fewer binders to convert the resulting pattern into force. This trade-off is strongest for mixed 1:4 particles, whose frequent entry into a locally depleted state can make cleavage reduce diffusivity.

## Cleaved trails promote detachment from receptor surfaces

We next confine the same bind–cleave model to a uniform 2D receptor surface. In Fig. 3a,b. trajectories begin from a common origin and stop when the particle loses attachment. Because the initial receptor landscape is uniform, their different exploration ranges arise entirely from the trails they write. In 3D, the probability of revisiting an older trail falls with time, but on a 2D surface a wandering particle returns to previously visited neighborhoods with probability one [19]. Polarized particles spread their narrow trails over a larger area, whereas mixed particles clear a body-width lane and cross it again sooner. Successive encounters with these cleaved regions leave fewer intact receptors for rebinding, making detachment inevitable (SI Section 3).

**Fig. 3.**
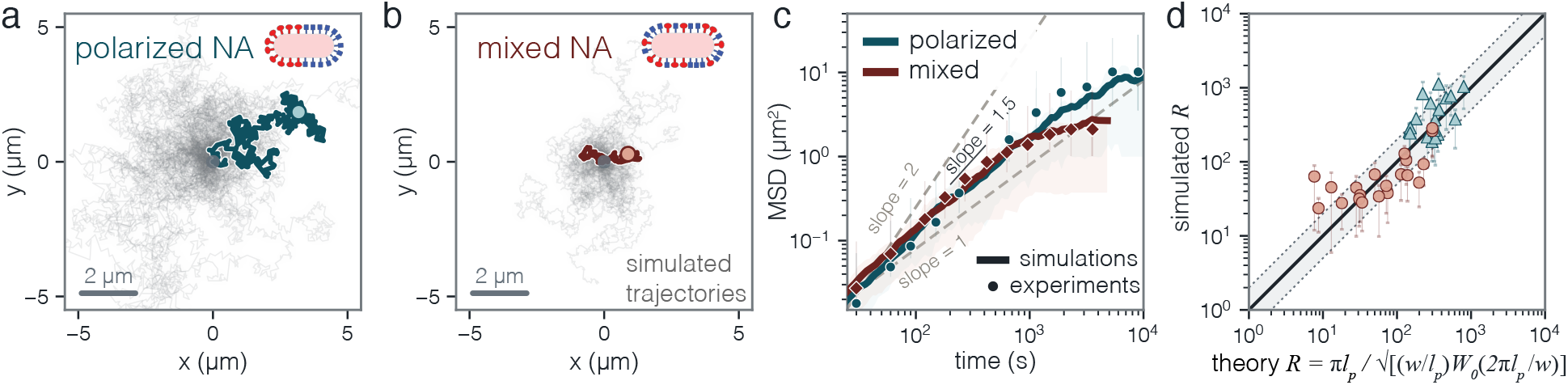
Repeated returns to cleaved trails set the exploration range before detachment. **a,b**, Simulated trajectories for polarized (**a**) and mixed (**b**) particles on a uniform 2D receptor surface, with one trajectory highlighted in each panel. **c**, Mean-squared displacements (MSDs) calculated from previously published wild-type (WT) and NA cytoplasmic-tail-deletion NAΔCT IAV trajectories (points) and from the corresponding polarized and mixed simulations (lines and interquartile envelopes). **d**, Mean exploration range at detachment from simulations versus the mean-field prediction. The solid line marks equality, the dotted lines mark a factor of two and error bars show interquartile ranges across trajectories.

We compared MSDs from previously published IAV surface trajectories with the corresponding polarized and mixed simulations (Fig. 3c) [13]. Both experiment and simulation initially approach ⟨Δ*r*^2^⟩∼ *t*^3*/*2^ before separating as returns to older trails accumulate. The simulations capture this common early spreading and later architecture-dependent detachment, using the architecture and NA-intensity mapping described in SI Section 3.

We define the exploration range *R* as the mean distance from the starting point when a particle detaches from the surface. It is set by the persistence length *ℓ*_*p*_, the distance covered before direction is lost, and the effective trail width *w*, the transverse span over which recrossing prevents renewed attachment. Their ratio compares fresh surface reached during a run with the reach of an old trail. Because each run adds a trail while the probability of crossing an existing segment decays only inversely with its age, encounters with trails of different ages accumulate. Summing over those trails gives, for *w* ≪ *ℓ*_*p*_,

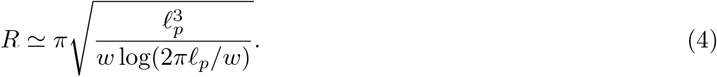

Persistence enters more strongly because longer runs carry the particle farther and spread earlier trails over a larger area. With *ℓ*_*p*_ and *w* calculated from molecular parameters and ligand geometry, the full mean-field expression captures the simulated ranges without fitting the range data (Fig. 3d; SI Section 3). Dimensionality therefore changes the outcome of the same particle–environment interaction: local receptor depletion controls 3D transport, whereas accumulated trail history fixes a finite surface exploration range before detachment.

## Receptor cleavage amplifies gradient guidance

We next impose the simplest environmental heterogeneity in Fig. 1. a shallow 3D gradient in the relative receptor density *ρ*(*z*) (Fig. 4a). Across its length *L*, the particle samples the fractional contrast Δ_*ρ*_ ≃ *L*∂_*z*_ log *ρ*. Because the gradient changes little over one particle length, the uniform theory gives the local speed, persistence, diffusivity and fraction of time a mixed particle remains mobile at *ρ*(*z*). The contrast Δ_*ρ*_ sets the bias toward denser receptors (SI Section 4).

**Fig. 4.**
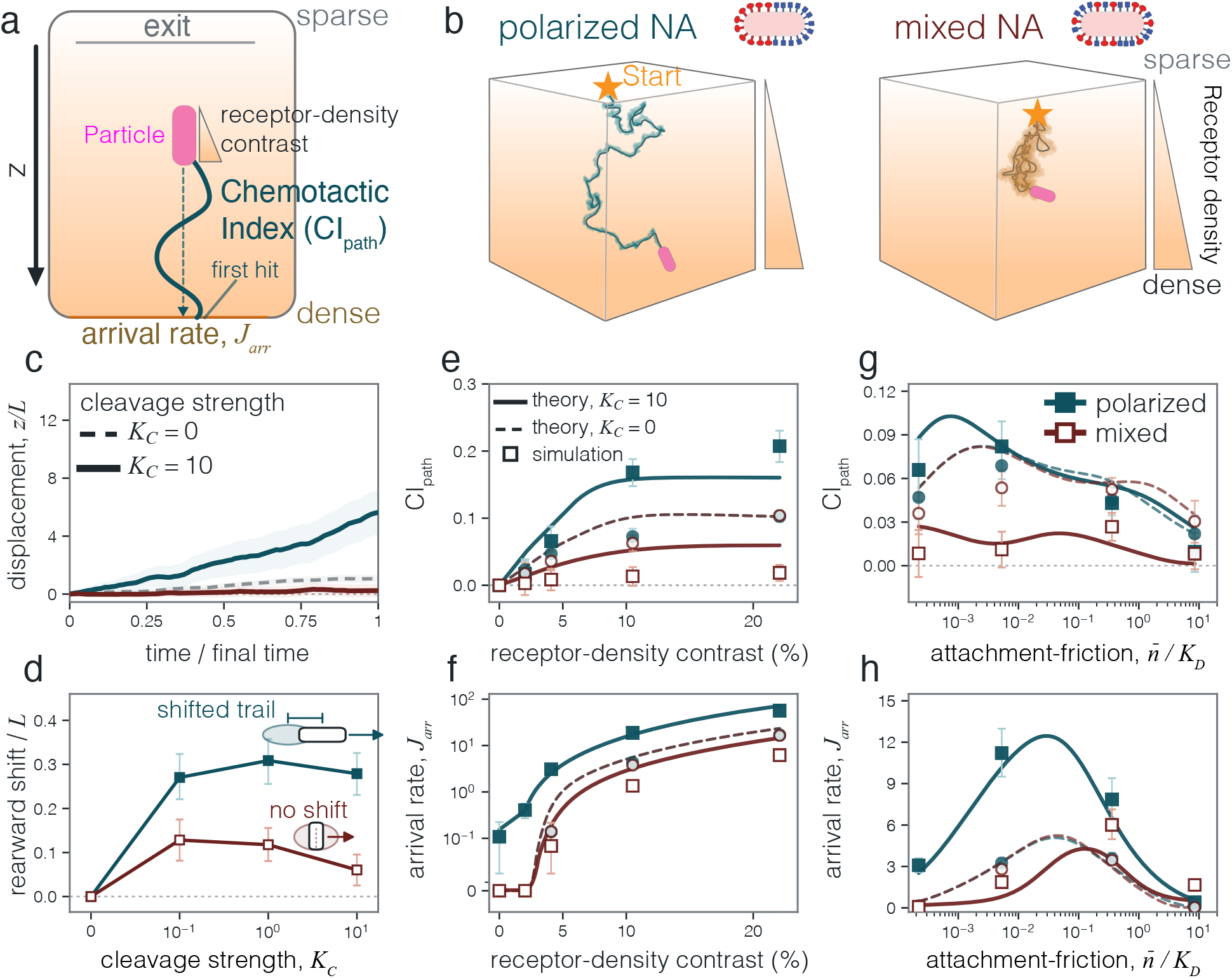
Polarized receptor cleavage amplifies guidance along a receptor-density gradient. **a**, Three-dimensional gradient geometry and definitions of the particle-scale receptor contrast and guidance measures. **b**, Representative polarized and mixed trajectories at *K*_*D*_ = 100; orange marks cleaved receptors. **c**,**d**, Particle displacement toward denser receptors (**c**) and displacement of the cleaved receptor region behind the particle (**d**) with increasing *K*_*C*_ . **e**,**f**, Pathwise chemotactic index CI_path_ (**e**) and normalized arrival rate at the denser boundary (**f** ) versus the receptor-density contrast across the particle at *K*_*D*_ = 500. **g**,**h**, The same observables versus 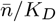, a measure of attachment-induced friction evaluated at the initial receptor density. Teal and red denote polarized and mixed particles; squares and solid curves show *K*_*C*_ = 10, while circles and dashed curves show *K*_*C*_ = 0. Symbols show simulations, curves show mean-field predictions and error bars are 95% trajectory-bootstrap intervals.

Binders nearer the higher-density side of the particle attach more often, allowing the particle to compare receptor abundance across its length [20, 21]. Reversible binding therefore biases both architectures toward denser receptors, and cleavage determines how that bias couples to the self-generated motion described above. Polarized particles move toward increasing density while the cleaved region shifts behind their binder-rich end, and both displacements grow with *K*_*C*_ (Fig. 4b–d). Mixed particles read and write within the same footprint, progressively removing the intact receptors needed to sustain motion. By displacing writing behind reading, the polarized architecture instead steepens the receptor difference sampled by the binders. Receptor destruction can therefore sharpen the directional cue.

At relative receptor density *ρ*, each binder samples an effective number *νρ* of receptors, giving 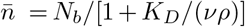 attached binders on average. The gradient of the multivalent binding free energy produces a binding-driven velocity toward denser receptors,

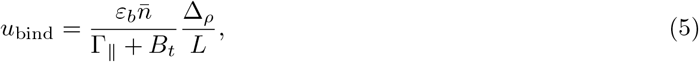

where *ε*_*b*_ is the energy scale of one attachment, *B*_*t*_ is the attachment-induced friction, and Γ_∥_ is the background friction along the particle axis. The effect of architecture is captured by *η*, the normalized difference in attachment probability between the ends facing higher and lower receptor density. The resulting velocities along the gradient are

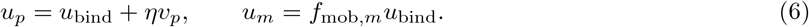

Here *v*_*p*_ is the local polarized run speed from the uniform theory. The larger attachment probability at the denser end favors binder-first runs, giving the mean projection *ηv*_*p*_. Mixed cleavage-driven runs are equally likely to point toward higher or lower receptor density and give no net motion along the gradient. Local receptor depletion further reduces *f*_mob,*m*_, the fraction of time the particle remains mobile and coupled to intact receptors.

We quantify directional progress using the pathwise chemotactic index CI_path_ = ⟨Δ*z*_*i*_*/*ℒ_*i*_⟩, where Δ*z*_*i*_ is the net displacement toward denser receptors and L_*i*_ is the 3D path length. Transport is measured by *J*_arr_, the normalized rate at which trajectories first reach the denser boundary. Here “chemotactic” denotes geometric alignment with the receptor gradient and does not imply internal signaling. Both increase with Δ_*ρ*_ (Fig. 4e,f). Without cleavage, polarized and mixed particles respond identically through reversible binding. Cleavage adds *ηv*_*p*_ for polarized particles but reduces the mixed response through *f*_mob,*m*_. At zero contrast, symmetry fixes CI_path_ = 0, but diffusion retains a finite arrival rate. Mean-field predictions use the molecular parameters and uniform-environment diffusivities without fitting the gradient simulations.

Varying affinity separates the optimum for guidance from that for rapid arrival (Fig. 4g,h). Because each attachment persists for 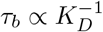, the friction from attached binders scales as 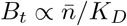. Weak attachment transmits little of the receptor-density difference, whereas near saturation the two ends respond similarly and persistent attachments slow the particle. Binding-driven motion therefore peaks at intermediate 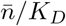. Within this range, CI favors weaker attachment because wandering adds path length without comparable up-gradient progress, while long-time diffusion can hasten first arrival at the denser boundary. The arrival optimum consequently lies at larger 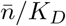 than the CI optimum (SI Section 4). Spatially displaced cleavage further strengthens polarized motion toward denser receptors. Together with the uniform-environment diffusivity, the resulting local drift predicts transport through receptor landscapes that vary slowly over one particle length.

## Local receptor gradients guide IAV transport

The gradient theory predicts a binding-driven bias toward denser receptors even without cleavage, while NA can alter both this bias and the mobility needed for it to accumulate into arrival. Previous studies have linked IAV trajectories on cell surfaces and receptor-bearing interfaces to receptor-rich domains and HA–NA activity [13, 14, 22, 23]. We tested this in a planar assay that placed a stable, reconstituted heterogeneous receptor landscape and viral trajectories in the same imaging plane, allowing us to compare each displacement with its local gradient and determine whether the resulting biases accumulated into arrival.

We crosslinked fluorescent sialylated CEACAM5 on supported lipid bilayers to create local heterogeneity, then imaged the receptor landscape and individual virions by three-color TIRF microscopy (Fig. 5a,b). For each displacement, *θ* was the angle between that displacement and the local receptor gradient (Fig. 5c), and CI_step_ = ⟨cos *θ*⟩ measured local geometric alignment, with positive values indicating steps toward denser receptors. Unlike the arrival rate at the denser boundary in the 3D simulations, *J*_net_ is signed, with first arrival on the denser side contributing positively and first arrival on the sparser side negatively. NA inhibition (NAI) with oseltamivir then separated local alignment from cleavage-dependent transport (SI Section 5).

**Fig. 5.**
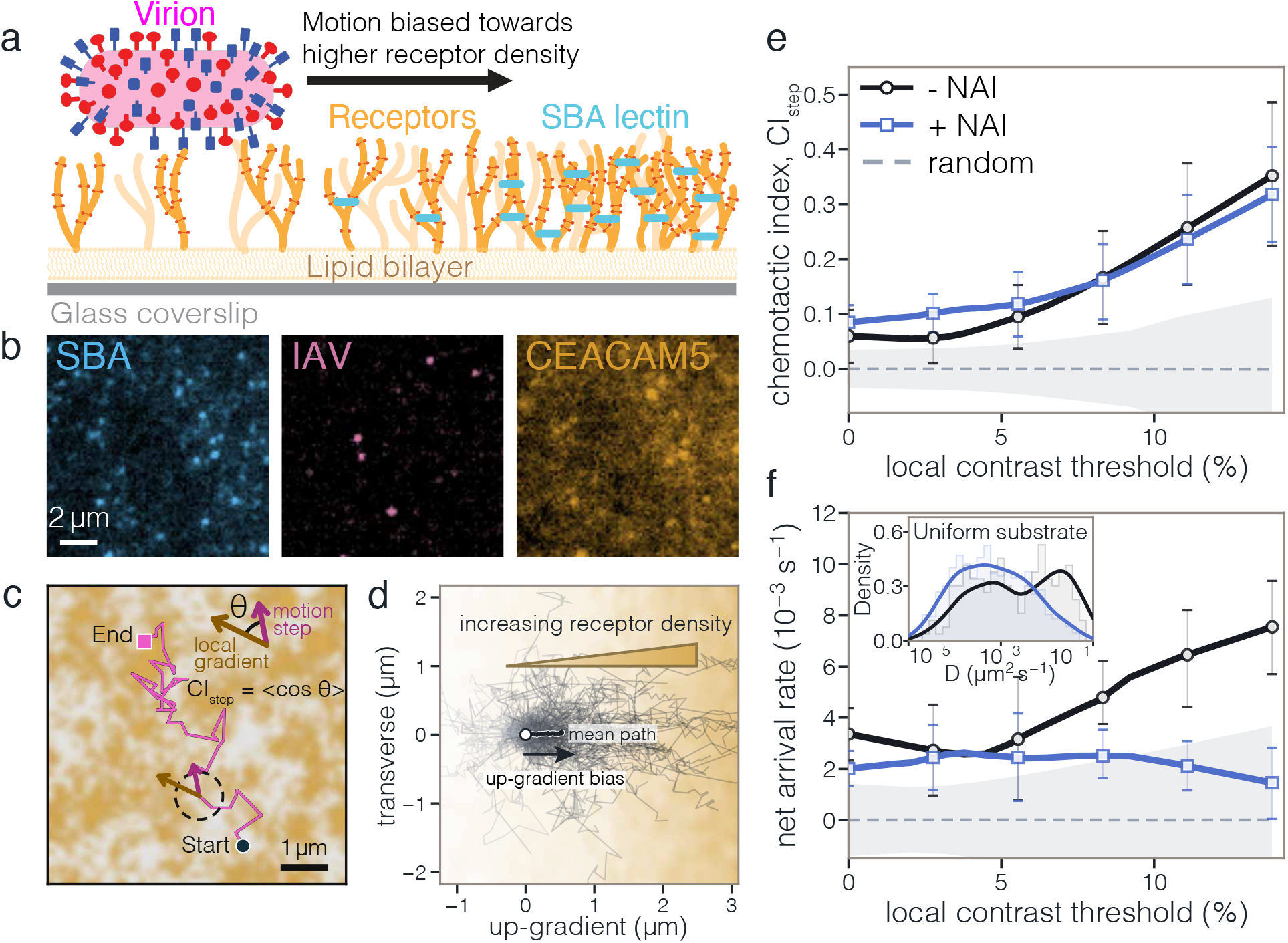
IAV motion aligns with local receptor gradients, while NA promotes arrival at denser regions. **a**, Heterogeneous receptor landscapes formed by soybean agglutinin (SBA) crosslinking of sialylated CEACAM5 on supported lipid bilayers. **b**, Matched three-color total internal reflection fluorescence (TIRF) images of SBA (488 nm), virions (561 nm) and receptors (647 nm). **c**, Representative trajectory over the measured receptor landscape, showing the local gradient and displacement angle *θ*. **d**, Trajectory fragments rotated with positive *x* along the local receptor gradient; the black curve shows the mean path. **e**, Particle-scale alignment CI_step_ versus the minimum receptor-contrast threshold; the gray band shows the randomized-direction null. **f**, Net arrival rate *J*_net_ for a 0.75 *µ*m arrival distance. Inset, diffusivity distributions on control bilayers without SBA crosslinking. Points and error bars in **e**,**f** show mean ± s.d. across three biological replicates. In condition-resolved curves, black denotes untreated particles (−NAI), and indigo blue denotes particles treated with the NA inhibitor oseltamivir (+NAI).

After trajectory fragments were rotated into their local gradient frames, their endpoints and mean path shifted toward the denser side (Fig. 5d). Alignment strengthened with local receptor contrast (Fig. 5e). Among the strongest 10% of receptor contrasts, both untreated and NA-inhibited particles aligned above the randomized-direction null, whereas their difference was not resolved across the three biological replicates. The retained alignment is consistent with the reversible-binding response predicted by the theory.

NA activity had a stronger effect on the accumulation of local biases into arrival. *J*_net_ increased with receptor contrast for untreated particles but was suppressed after NA inhibition (Fig. 5f). Within the same contrast range, NA inhibition reduced *J*_net_ approximately threefold, with the same direction of change in all three replicates. On control bilayers without SBA crosslinking, NA inhibition also reduced the fast-diffusing component of the apparent-diffusivity distribution (Fig. 5f. inset).

Particle size, HA–NA abundance and surface organization were not resolved for individual virions, so trajectories cannot be assigned to the idealized architectures. Even across this heterogeneity, local alignment persisted after NA inhibition while net arrival fell, showing that reversible binding generates local alignment, whereas NA-dependent mobility accumulates successive biases into transport.

## IAV heterogeneity spans distinct transport states

The broad diffusivity distribution in Fig. 5f (inset), including its NA-sensitive fast component, shows that the tracked virions span distinct transport states. Previous measurements in other IAV strains reveal similarly broad physical variation, with HA and NA abundances spanning two orders of magnitude and particles shorter than 300 nm occupying broad HA:NA modes near 2:1 and 10:1 [18]. Low-fidelity assembly generates this variation, while host-cell conditions and external pressures reshape the resulting phenotype distribution [18, 24, 25]. Because binding and cleavage enter transport nonlinearly, the distribution over transport states—not the mean composition—is the relevant physical object.

Along the route to infection, mucus, adhesive decoys and receptor-rich cell surfaces place competing demands on the same HA–NA machinery [26–28]. Continued motion enables *exploration*, cleavage permits *escape* from adhesive decoys, and high-avidity *exploit* states sustain attachment to receptor-rich surfaces. Stronger binding increases receptor coupling but slows transport, whereas cleavage can free an attached particle or destroy the receptors needed for continued motion. We use exploit only for this persistent pre-entry contact; cellular uptake and ensuing HA-driven membrane fusion lie outside the transport model [29].

We map these transport states onto a plane defined by the mean attached-binder fraction *ϕ*_*b*_, which measures collective avidity, and cleavage capacity *χ*_*C*_, which combines cleaver number and activity (Fig. 6a,b; SI Section 6). Increasing *ϕ*_*b*_ carries a particle from weak receptor sampling through mobile exploration to persistent attachment at low *χ*_*C*_. Increasing *χ*_*C*_ extends polarized exploration because cleavage is displaced from subsequent binding, whereas in mixed particles it can complete escape when only a few attachments remain or remove the local receptors supporting a strongly attached particle. Architecture can therefore direct the same intermediate-avidity state toward exploration or escape. Each color in Fig. 6 shows the best score for one function after optimizing binder allocation, so overlapping regions represent distinct particles at the same (*ϕ*_*b*_, *χ*_*C*_).

**Fig. 6.**
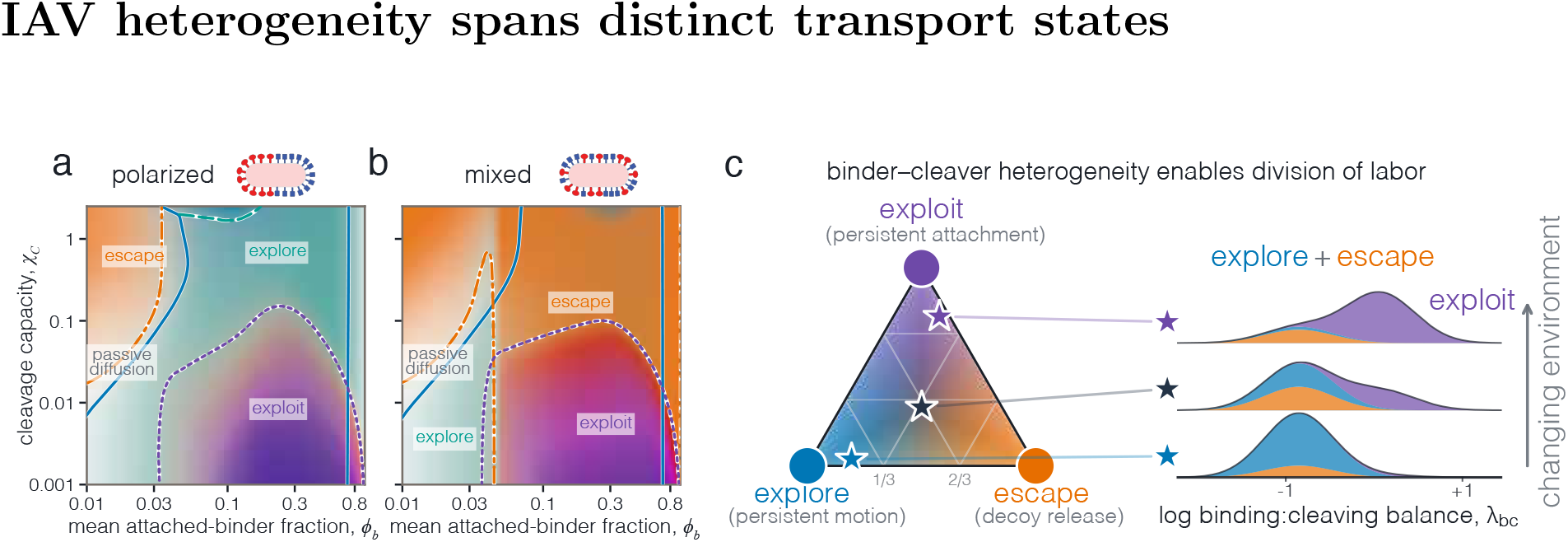
Binder–cleaver heterogeneity distributes particles across explore, escape and exploit states. **a**, Polarized architecture mapped by the mean attached-binder fraction, 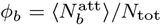, and cleavage capacity, *χ*_*C*_ = *K*_*C*_ *N*_*c*_*/N*_tot_. Colors show the strongest attainable function after optimizing binder allocation. **b**, Mixed architecture in the same coordinates. **c**, Population fractions that maximize the least-supplied function for exploit-biased, equal and explore-biased demands. Each adjacent density is the projection of the corresponding star onto the binder:cleaver balance *λ*_bc_. The stacked blue, orange and purple areas integrate to the explore, escape and exploit fractions, respectively, and the solid outline is their sum. Theory fixes the density centers; a common width only displays their overlap.

The population calculation in Fig. 6c draws polarized explorers, mixed escape specialists and exploit specialists from the same mean-field landscape and assigns their weights to maximize the least-covered function so that no single demand becomes a bottleneck. Moving from exploration-biased to exploit-biased demands transfers weight from polarized explorers to binder-rich exploit states while retaining a mixed escape component. These transport phenotypes show why a broad distribution across separated states can accommodate changing landscape demands more effectively than a narrow distribution around the mean. Assembly pressures can redistribute particles among these states, while NA inhibition (Fig. 5f. inset) moves states toward lower *χ*_*C*_. A self-edited receptor landscape can thus convert particle heterogeneity into complementary transport functions through independent particle–landscape feedback.

## Discussion

Our results place activity in the coupled dynamics of a particle and a writable landscape, where reversible binding reads receptor availability, irreversible cleavage writes a persistent change and rebinding biases the next fluctuation-driven displacement. The free-energy drop due to receptor cleavage provides the nonequilibrium drive without ATP hydrolysis or a molecular motor. Avidity and architecture determine the particle’s response to the edited landscape and thereby set its diffusivity. The coupled particle–landscape system is therefore the active unit.

Irreversible substrate modification rectifies fluctuations in burnt-bridge ratchets, DNA walkers, collagenases and synthetic protease motors [9, 10, 30–32]. Extending this principle from a 1D track to higher dimensions makes re-encounter probability a control variable. At the level of the particle trajectory, the resulting feedback produces a self-interacting, non-Markovian walker whose exploration depends on its own footprints [7, 33]. Old trails become less likely to be revisited in 3D, whereas in 2D the particle repeatedly crosses the path it has already written. The normalized exploration range *R/ℓ*_*p*_ then collapses onto *ℓ*_*p*_*/w*, yielding 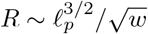. The same chemistry can enhance transport in 3D yet limit 2D exploration to a finite range.

A receptor gradient turns this coupling into guidance. Multivalent binding can drive interfacial motion along gradients of receptor availability [34], while spatially displaced cleavage sharpens the contrast that binders later sample. Cue consumption can guide motile cells by reshaping diffusible fields [35], and degradation can increase directional information when coupled to diffusion [36]. In IAV, receptor readout and editing occur within the same nanoscale contact zone, where binder–cleaver architecture reshapes the receptor contrast that generates force. Simultaneous imaging of individual virions and the fluorescent receptor landscape shows that IAV motion is biased up local receptor gradients. NA inhibition preserves alignment but strongly reduces net arrival. Particle guidance can thus emerge without internal signaling.

Nonlinear transport makes the phenotype distribution more informative than its mean. Receptor landscapes map different phenotypes onto exploration, escape and exploit, providing a physical mechanism through which heterogeneity could hedge against changing environments. Whether viral evolution has tuned the distribution in this way remains open [37, 38].

Bind–cleave chemistry is one realization of read–write feedback. Writing may add, remove or transform the local environment, with the resulting dynamics set by avidity, the lifetime of the change, reader–writer organization and dimensionality. Information thermodynamics links environmental memory to predictive content and energetic cost [39–41]. For such systems, directional information gained per environmental modification may be a useful performance metric. Across scales, cells can consume and reshape the cues they follow [35], extracellular vesicles can carry adhesive integrins or matrix-remodeling proteases, and compositionally tunable lipid nanoparticles offer a synthetic route [42–44]. Intelligent-matter approaches integrate sensing, actuation and memory within a material [45]; read–write feedback instead distributes these capabilities across a moving object and its surroundings. A writable landscape shared by many cells or particles could turn individual modifications into collective memory and mediate indirect coordination [46]. Functions associated with living systems—active transport, guidance and memory—can therefore emerge from feedback between a moving object and a writable environment.

## Methods

### Stochastic receptor-editing simulations

We represented the particle as a rigid rod of length *L*, with *N*_tot_ = int(*L*) ligand sites along its axis. For *L* = 20, this gives 20 sites at unit spacing. Of these, *N*_*b*_ were binders and *N*_*c*_ = *N*_tot_ − *N*_*b*_ were cleavers, arranged in the polarized or mixed geometries defined in the main text. In polarized particles, the two ligand types occupied separate axial blocks. In the mixed layout, cleavers occupied both endpoints and the remaining cleavers were distributed approximately uniformly among the interior sites. With particle center **X**, axis **u** and fixed axial coordinates *s*_*i*_, the ligand positions were **r**_*i*_ = **X** + *s*_*i*_**u**. This construction averaged over positions around the rod circumference while preserving axial organization. Uniform 3D receptors occupied a simple cubic lattice of spacing *d*_rec_. Receptors were fixed during each trajectory, as appropriate when redistribution is slower than rebinding or return to a trail. Cleaved-receptor trails have also been observed in native mucus, and directional bias persists with slow receptor diffusion and replenishment [13]. Further details are given in SI Section 1. Each receptor was free, binder-bound or irreversibly cleaved. We measured time in units of the inverse binding-rate prefactor 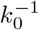. With the Gaussian interaction weight *ω*(*r*) = exp(− *r*^2^*/α*^2^) and interaction range *α*, the reaction rates were

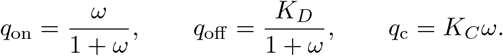

Thus *K*_*D*_ and *K*_*C*_ are the dimensionless unbinding and cleavage parameters. Dimensional rates are recovered by multiplying by *k*_0_, and *q*_on_*/q*_off_ = *ω/K*_*D*_ sets the equilibrium binding weight in the mean-field theory. An attached binder acted as a zero-rest-length harmonic spring of stiffness *k*_*s*_, supplying force −*k*_*s*_***δ*** and torque about the particle center, where ***δ*** is the binder–receptor displacement. Between reactions, the particle followed overdamped translation and rotation,

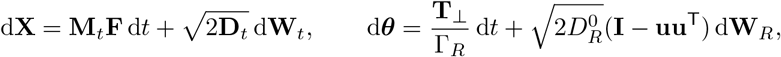

where **F** and **T**_⊥_ are the total receptor force and transverse torque, d***θ*** is the rotation vector, and d**W**_*t*_ and d**W**_*R*_ are independent Wiener increments. The tensors **M**_*t*_ and **D**_*t*_ contain the axial and transverse mobilities and diffusivities. We set *k*_*s*_ = 1 and define the attachment-work scale *ε*_*b*_ = *k*_*s*_*α*^2^*/*2 = 0.125. Using

*ε*_*b*_ as a reference energy scale gives 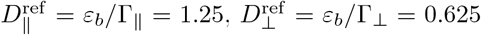 and 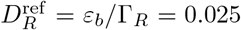, with Γ_∥_ = 0.1, Γ_⊥_ = 0.2 and Γ_*R*_ = 5. The background diffusivities were 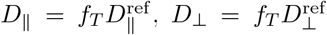 and 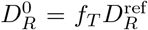. The dimensionless factor *f*_*T*_ scales background translational and rotational fluctuations without changing receptor forces or reaction rates. Uniform 3D and gradient simulations used *f*_*T*_ = 0.01, giving *D*_∥_ = 0.0125, *D*_⊥_ = 0.00625 and 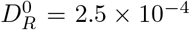. Surface-return simulations used *f*_*T*_ = 0, leaving motion driven by receptor binding, unbinding and cleavage.

For total reaction rate *Q* = ∑_*j*_ *q*_*j*_, we drew the waiting time as − log *U/Q*, where *U* is uniform on (0, 1), and chose reaction *j* with probability *q*_*j*_*/Q*. Between reactions, we updated particle position, orientation and reaction rates with adaptive time steps. One simulation coordinate unit represents 10 nm. Thus *L* = 20 represents 200 nm and *α* = 0.5 represents 5 nm. SI Section 1 details the reaction–motion algorithm and numerical checks.

### Transport measurements

We measured body-axis alignment as 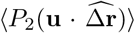, where *P*_2_(*x*) = (3*x*^2^ − 1)*/*2 is the second Legendre polynomial. Persistence came from the 1*/e* crossing of ⟨**u**(*t*) · **u**(0)⟩ for polarized particles and ⟨*P*_2_[**u**(*t*) · **u**(0)]⟩ for mixed particles. Polarized speed was the median early axial velocity. For mixed particles, we estimated speed from correlations between successive transverse velocities, after subtracting the matched *K*_*C*_ = 0 correlation and accounting for directional decay. We excluded later intervals in which no intact receptor was accessible. Effective diffusion came from late-time ensemble MSD slopes, 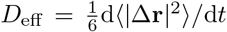, and Δ*D*_eff_ was calculated relative to the matched *K*_*C*_ = 0 condition. Error bars show trajectory interquartile ranges for *P*_2_, 5–95% trajectory-bootstrap intervals for *v* and *τ*, 95% paired ensemble-MSD intervals in Fig. 2f and interquartile paired ensemble-MSD intervals in Fig. 2g. SI Section 2 describes how these quantities and uncertainty intervals were calculated.

### Mean-field calculation

We assumed that binding reached local equilibrium, that bonds formed and broke over distances shorter than those over which receptor density changed, and that each run extended beyond one interaction range. The calculation links the friction created by attached binders and rotational persistence to receptor contrast, run statistics and diffusion (SI Section 2). Two Gaussian-weighted averages give

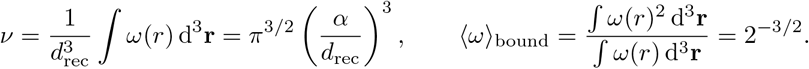

The free state has weight one, while all attached configurations have combined weight *ν/K*_*D*_, giving the probability *p*_*b*_ = *ν/*(*K*_*D*_+*ν*) that one binder is attached and the average number 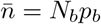 of attached binders. Weighting the local lifetime (1 + *ω*)*/K*_*D*_ by the distribution of attached configurations gives *τ*_*b*_ = *κ*_*b*_*/K*_*D*_, where *κ*_*b*_ = 1 + 2^−3*/*2^. Independent binding and unbinding events then give

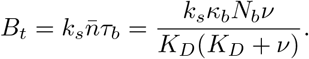

Binding equilibrium controls both collective attachment strength (avidity) and the friction generated as bonds form and break. Eliminating *K*_*D*_ in favor of 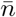 gives

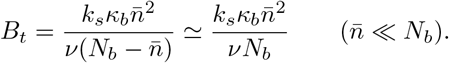

The probability of at least one attachment and the fraction of the unbound mobility retained after binding are

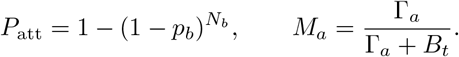

Small *P*_att_ limits force transmission, while attachment-induced friction lowers *M*_*a*_. These are the two factors plotted in Fig. 2i.

As the particle passes, cleavage leaves a local intact-receptor fraction *f*_int_ = exp(−Ω_*a*_*/v*_*a*_), where Ω_*a*_ follows from *K*_*C*_, cleaver number and ligand positions. Binders convert two sampled receptor fractions into the free-energy contrast

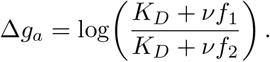

The two receptor fractions sampled by the binders are

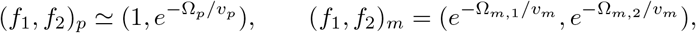

where Ω_*m*,1_ ≠ Ω_*m*,2_ reflect how mixed cleavers sample the two sides. The polarized contrast saturates because one sampled region remains intact, whereas the mixed contrast vanishes when both regions are intact or depleted. Dividing *ε*_*b*_Δ*g*_*a*_ by the sampling distance and averaging over ligand geometry gives *F*_*a*_ in Eq. 2. We obtained the architecture-specific Ω_*a*_ and *F*_*a*_ by averaging the Gaussian interaction weights along the ligand paths. At weak cleavage, *f*_int_ ≃ 1 − Ω_*a*_*/v*_*a*_ and *F*_*a*_ ∝ *K*_*C*_*/v*_*a*_. Inserting this expansion into Eq. 2 gives 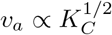. The sign of ⟨*P*_2_⟩ selects the background friction in the force balance. We used Γ_*p*_ = Γ_∥_ for polarized axial motion and Γ_*m*_ = Γ_⊥_ for mixed transverse motion.

Each attached binder also produces rotational friction proportional to *k*_*s*_*p*_*b*_*τ*_*b*_ times its squared lever arm. For the symmetric mixed layout,

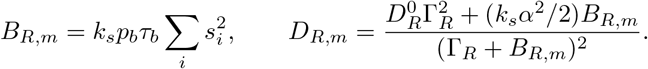

The polarized calculation includes translation–rotation coupling, but both architectures approach the limits in Eq. 1. For long-lived attachments, transverse relaxation makes rotational persistence depend on the squared spread of the binders about their mean position, 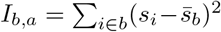, where 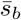 is the mean binder coordinate. The head-to-tail orientation (*ℓ* = 1) of a polarized particle decays three times more slowly than the axis-only orientation (*ℓ* = 2) of a mixed particle, giving 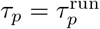 and 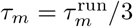. These orientational timescales enter the velocity-correlation time *τ*_*v,a*_, together with any loss of directed motion before the particle reorients. The latter matters for mixed particles that cross their body-wide depleted region (SI Section 2). The receptor-driven contribution then follows from the Green–Kubo relation

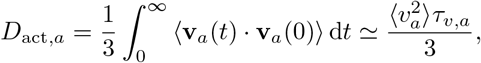

where the final form uses a single dominant correlation time.

A mixed particle that reverses can cross the recently cleared region of width *w*_*m*_ = *L* + 2*α*. The entry rate *k*_in_ is the probability of crossing this region and losing all receptor attachments, divided by the directional lifetime. Transverse diffusion gives 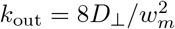, and balancing these rates gives *f*_mob,*m*_ = *k*_out_*/*(*k*_in_+*k*_out_). Polarized particles remain coupled to intact receptors, so *f*_mob,*p*_ ≃ 1. With *D*_bu,*a*_ = *D*_eff,*a*_(0), the diffusivity shift for either architecture is

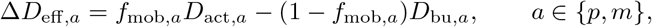

as in Eq. 3. Figure 2h compares the difference in remaining receptor fraction *C*_*a*_ = 1 − *f*_low_*/f*_high_, where *f*_high_ and *f*_low_ are the larger and smaller sampled fractions, with the locally depleted fraction 1 − *f*_mob,*m*_, which is zero for polarized particles. The normalized curves cross when receptor contrast and local depletion become comparable, although the diffusivity maximum also depends on speed and persistence. We included only the most recent depleted region because a 3D walk returns only transiently to older bounded regions. We varied the binder:cleaver ratio while keeping *N*_tot_ fixed. Changing *N*_*b*_ altered 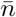, *B*_*t*_ and *P*_att_, whereas changing *N*_*c*_ altered Ω_*a*_. All other parameters were fixed.

### Surface-return simulations and mean-field analysis

Surface simulations used the same stochastic reaction rules and polarized or mixed ligand arrangements, with receptors confined to one plane. The main data in Fig. 3 used *K*_*D*_ = 10 and *f*_*T*_ = 0 to isolate repeated encounters with the cleaved trail. At this affinity, the uniform 3D diffusivity shift remained positive and showed no measurable dependence on *f*_*T*_ over the sampled range (Supplementary Fig. S1). A trajectory ended when loss of its final attachment left no intact receptor within reach. We calculated MSDs from previously published experimental trajectories and the matched simulations (Fig. 3c). WT used the polarized architecture, whereas NAΔCT used the mixed architecture and a tenfold reduction in *K*_*C*_ set by the measured NA intensity. All other kinetic parameters were fixed. We converted simulation units using 10 nm per length unit and 0.09 s per time unit, corresponding to an effective *k*_0_ = 11.1 s^−1^. To include all particles in the late-time MSD, we held their positions fixed after detachment. The exploration range was each particle’s final distance from its starting point. Figure 3d reports the mean range *R*.

While a particle remains attached, repeated binding and unbinding produce diffusive motion, to which persistent runs add. The resulting diffusivity and persistence length are 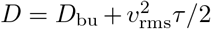 and 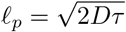, respectively, where *v*_rms_ is the root-mean-square run speed and *D*_bu_ is the diffusion produced by stochastic binding and unbinding. The effective cleaved width is

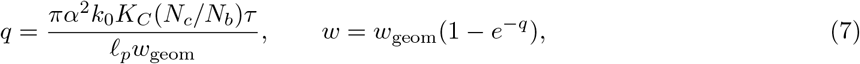

where *q* is the mean cleavage accumulated across the trail during one run, *w*_geom_ = 2*α* for polarized particles and *w*_geom_ = *L*_*c*_ + 2*α* for mixed particles, and *L*_*c*_ is the axial distance between the outermost cleavers. Thus *w* is the effective transverse span over which enough receptors have been removed to end attachment on a later crossing. We include *k*_0_ because *τ* is expressed in dimensional time. We obtained the range by summing return probabilities over trails of different ages. After *j* further runs, the probability of crossing one old trail is *w/*[2*πℓ*_*p*_(*j* + 1)]. Because each run adds another trail, the cumulative crossing probabilities after *n* completed runs sum to

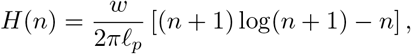

and the surviving attached fraction is *e*^−*H*(*n*)^. Setting *H*(*n*_∗_) ≃ 1 gives

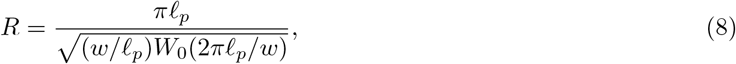

where *W*_0_ is the principal branch of the Lambert *W* function [47, § 4.13]. SI Section 3 gives the finite-time correction and comparison with the simulated exploration ranges. In the narrow-trail limit, *W*_0_(2*πℓ*_*p*_*/w*) ≃ log(2*πℓ*_*p*_*/w*), recovering

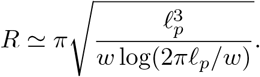

### Linear-gradient simulations and mean-field analysis

Gradient simulations used the same reaction rules and mechanical parameters, with all three background diffusion coefficients scaled by *f*_*T*_ = 0.01. SI Section 4 describes receptor placement and boundary conditions. The spacing between successive receptor planes varied geometrically along one Cartesian axis, while the two transverse spacings remained fixed. Orienting *z* toward the dense side gives the local relative spacing *s*_*ρ*_(*z*) = 1 − *βz*, where *β* sets the gradient strength, and density *ρ*(*z*) = 1*/s*_*ρ*_(*z*) before the imposed spacing limits are reached. The conditions *βα* ≪ 1 and *βL* ≪ 1 kept the density nearly uniform over the interaction range and particle length. The particle-scale contrast was

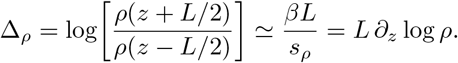

At local density *ρ*, the statistical weight for one binder is *Z*_*b*_ = 1 + *νρ/K*_*D*_, binding probability *p*_*b*_(*ρ*) = *νρ/*(*K*_*D*_ + *νρ*) and average number of attached binders 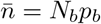. The Gaussian interaction and spring give the attachment energy *ε*_*b*_ = *k*_*s*_*α*^2^*/*2. The local multivalent binding free energy is

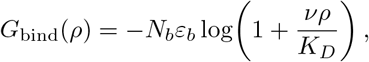

whose spatial derivative gives 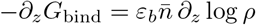. Equivalently, with *ρ*_±_ = *ρ*(*z ± L/*2), the binding work across the particle is

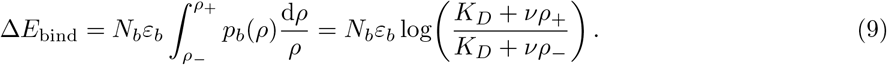

In the shallow-gradient limit, both forms give 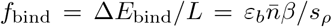. The average lifetime of one binder–receptor attachment is *τ*_*b*_ = *κ*_*b*_*/K*_*D*_, where *κ*_*b*_ = 1 + 2^−3*/*2^ is the attachment-lifetime factor defined above, and 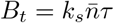_*b*_. Force balance gives

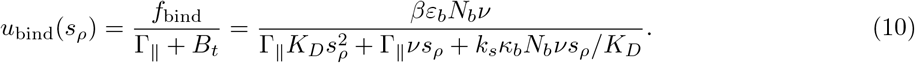

Thus 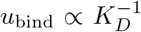 when binding is weak, while *u*_bind_ ∝ *K*_*D*_ when attachment-induced friction dominates. Because 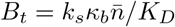, the plotted ratio 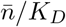 measures attachment-induced friction. The limiting forms are

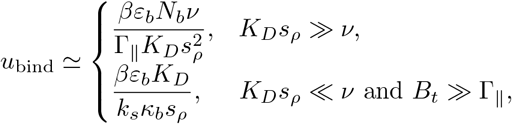

and their crossover occurs at

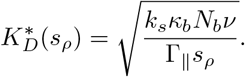

When few binders are attached, 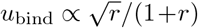, with *r* = *B*_*t*_*/*Γ_∥_, and peaks when attachment-induced and background friction are equal. To obtain the plotted curves, we averaged the product of force and mobility over the binomial distribution of the number of attached binders.

With *p*_*b*,±_ = *p*_*b*_(*ρ*_±_), the normalized difference in attachment probability between the ends facing higher and lower receptor density is

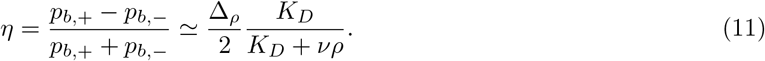

When few binders are attached, *η* approaches Δ_*ρ*_*/*2. As binding saturates, the difference in attachment probability between the two ends vanishes. For polarized particles, *η* multiplies the local run speed *v*_*p*_(*ρ*) from the uniform theory. Mixed particles run toward higher and lower receptor density with equal probability, so their cleavage-driven runs produce no net drift. Time spent locally depleted reduces their binding-driven drift by *f*_mob,*m*_(*ρ*), giving

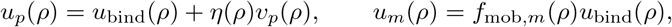

which is Eq. 6. Here *a* = *p, m* denotes the polarized or mixed architecture and *u*_*a*_ its local drift. From the uniform calculations, we took the diffusivity *D*_path,*a*_ over the sampling interval and the long-time diffusivity *D*_arr,*a*_ over the full boundary-crossing time.

For trajectory *i*, Δ*z*_*i*_ is the net displacement toward denser receptors, ℒ_*i*_ the 3D path length and *t*_hit,*I*_ the time of first arrival at a boundary. We calculated

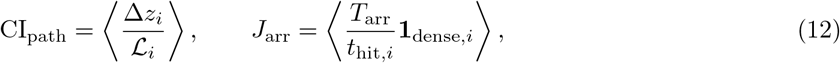

where *T*_arr_ is the observation time and **1**_dense,*i*_ = 1 only when the denser boundary was reached first within *T*_arr_. Trajectories that reached the sparse boundary first, or neither boundary, contributed zero.

Over a sampling interval Δ, drift contributes *u*_*a*_Δ. For weak drift, a 3D Gaussian step has mean length 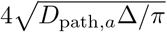. For arrival, we assumed that drift and diffusion changed little before the dense boundary was reached. The sparse boundary was much farther away on this timescale, and *T*_arr_ exceeded the typical arrival time. This gives

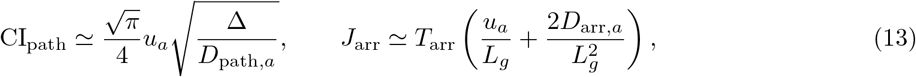

where *L*_*g*_ is the initial distance to the denser boundary. At zero drift, CI_path_ = 0, but diffusion still produces arrivals at the dense boundary. The affinity optima differ because *D*_path,*a*_, which measures wandering between recorded positions, lowers CI, whereas *D*_arr,*a*_ helps a particle reach the boundary. The CI optimum therefore occurs at smaller 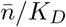 than the arrival optimum. For the plotted curves, drift and diffusion varied with *ρ*(*z*), both boundaries were absorbing and the simulation observation times were used. The full calculation also included the cleavage-driven motion of polarized particles, local receptor depletion around mixed particles and diffusion to both boundaries.

### Supported lipid bilayer functionalization

No. 1.5H glass coverslips were sonicated in 50% (v/v) isopropanol for 15 min, rinsed twice in Milli-Q water for 5 min each and treated for 3 min with freshly prepared 3:1 (v/v) sulfuric acid and 30% hydrogen peroxide. After thorough rinsing, small unilamellar vesicles composed of DOPC with 1 mol% 18:1 DGS-NTA(Ni) (Avanti Polar Lipids, catalog no. 790404) were deposited on the cleaned coverslips to form supported lipid bilayers (SLBs). Subsequent functionalization and washing used HEPES-buffered saline (HBS; 25 mM HEPES, 150 mM NaCl, pH 7.4). SLBs were blocked with 0.1% (w/v) BSA in HBS, washed three times and incubated for 15 min with 25 nM AF647-labeled human CEACAM5 bearing a C-terminal polyhistidine tag (Sino Biological, catalog no. 11077-H86H-SG), thereby anchoring CEACAM5 to DGS-NTA(Ni). After three washes, the SLBs were incubated for 30 min with 80 *µ*g ml^−1^ fluorescein-conjugated soybean agglutinin (SBA; Vector Laboratories, catalog no. FL-1011-2) to generate heterogeneous receptor distributions, then washed three more times. SBA crosslinks GalNAc/galactose motifs while leaving terminal sialic-acid ligands available for IAV attachment [48, 49].

### Cell culture and IAV propagation

Madin–Darby canine kidney (MDCK) cells were obtained from the UC Berkeley Cell Culture Facility and routinely screened for mycoplasma contamination in the laboratory. Cells were seeded in 12-well plates to reach approximately 70% confluence on the day of infection. Mouse-adapted reassortant X-31b IAV (H3N2; BEI Resources, catalog no. NR-3483) was stored at −80 ^◦^C and rapidly thawed in a 37 ^◦^C water bath. After two washes with sterile phosphate-buffered saline (PBS), we added 100 *µ*l of virus stock per well for 1 h, with gentle agitation every 15 min. The inoculum was removed, cells were washed twice and each well received 500 *µ*l infection medium containing minimum essential medium, 0.25% (w/v) bovine serum albumin, 1% (v/v) penicillin–streptomycin, 2.2 mg ml^−1^ sodium bicarbonate and 1 *µ*g ml^−1^ TPCK-treated trypsin. Cells were incubated overnight at 37 ^◦^C. At 18 h post-infection, neuraminidase was added to 10 mU ml^−1^ for 2 h. Plates were then gently agitated to release surface-associated virions, and pooled supernatants were clarified at 2,000 × *g* for 5 min. The supernatant was divided into 100 *µ*l aliquots for immediate use or flash-frozen in liquid nitrogen and stored at −80 ^◦^C.

### IAV labeling, binding and TIRF microscopy

Thawed 100 *µ*l virus aliquots were incubated with 0.5 *µ*M octadecyl rhodamine B chloride (R18) for 1 h at room temperature. Capto Core 700 resin (1 ml) was equilibrated with HBS through three cycles of resuspension and settling, then divided into two equal portions. R18-labeled virus was passed through the two portions in sequence, and the virus-containing supernatant was collected after each portion settled. Labeled IAV (5 *µ*l) was added to each prepared SLB and incubated for 10 min at room temperature. The SLBs were washed three times with 100 *µ*l HBS to remove unbound particles. Coverslips were imaged by total internal reflection fluorescence (TIRF) microscopy on a Nikon Ti2 inverted microscope equipped with an iLas2 ring-illumination module (GATACA Systems), a Nikon Apo TIRF 100× oil-immersion objective (NA 1.49) and a Hamamatsu ORCA-Fusion sCMOS camera (C14440-20UP). Three-color image sets were acquired every 2 s with 61-ms exposures and 2 × 2 camera binning (effective pixel size, 0.13 *µ*m) using NIS-Elements AR 5.42.03. The 488-nm, 561-nm and 647-nm channels recorded SBA, IAV and CEACAM5, respectively. We imaged untreated samples (− NAI) and samples treated with 100 *µ*M oseltamivir (+NAI) in three biological replicates. Labeling, SLB preparation and all subsequent assay steps were repeated independently for each replicate.

### Single-virion tracking and gradient analysis

Particle positions were detected and linked using recording-specific noise estimates. SI Section 5 gives the tracking criteria and robustness tests. The initial receptor image *I*(**r**) was corrected for uneven illumination and Gaussian-smoothed with *σ*_*g*_ = 0.25 *µ*m. Its first angular harmonic on a ring of radius *r*_*g*_ = 0.5 *µ*m defined the local cue direction and fractional front-to-back contrast *C*_*i*_ = 4 **h**_*i*_ */Ī*_*i*_, where **h**_*i*_ is the first-harmonic vector and *Ī*_*i*_ is the mean receptor intensity on the ring. The chemotactic index CI_step_ = ⟨cos *θ*_*i*_⟩ was calculated from non-overlapping 0.5 *µ*m first-passage displacements. A shared threshold selecting the strongest 10% of receptor contrasts was set by pooling both treatments and all biological replicates, then applied unchanged. For *J*_net_, consecutive selected frame-to-frame displacements were projected onto the changing local cue. First arrival at ±0.75 *µ*m contributed ±1*/t*_hit_, and non-arrivals contributed zero. We averaged measurements first within tracks, then recordings and biological replicates. Biological replicates were the units of statistical comparison. On control membranes without SBA crosslinking, apparent diffusivity *D*_app_ came from time-averaged MSD fits. We fitted a two-component Gaussian mixture to untreated log_10_ *D*_app_ and used the same mixture to classify both treatments. We used paired biological-replicate means for treatment comparisons. Permutation tests shuffled treatment labels among recordings within each replicate, and randomized-direction nulls were averaged first within tracks, then recordings and replicates.

### Population transport-state calculation

A particle state is set by its architecture, binder number *N*_*b*_, cleaver number *N*_*c*_ = *N*_tot_ − *N*_*b*_, *K*_*D*_ and *K*_*C*_. We summarize binding and cleavage with

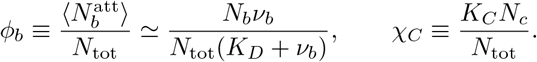

Here *ν*_*b*_ = *νρ*_ref_ is the number of receptors accessible to one binder at the reference density *ρ*_ref_ . The quantity *ϕ*_*b*_ combines binder number and affinity, while *χ*_*C*_ combines cleaver number and activity. From the uniform and gradient theories, we calculated the diffusivity gain, pathwise chemotactic index, probability of at least one attachment *P*_att_, cleavage probability *P*_cleave_, and *P*_local_, the probability that local cleavage removes the receptors supporting an existing attachment in a mixed particle. These probabilities combine as

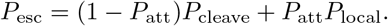

The first term describes cleavage-mediated escape when sustained attachment is already unlikely. The second describes escape after an attached particle clears its local receptor footprint. For polarized particles, *P*_local_ = 0 at this order. We normalized the positive diffusivity gain and pathwise chemotactic index to obtain spreading and sensing scores, then weighted both by 1 − *P*_esc_. The explore score is the larger of the two, and the escape score is *P*_esc_. The exploit score is

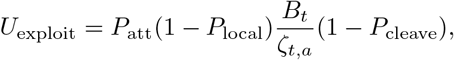

which combines persistent attachment, attachment-induced friction and preservation of nearby receptors. Here *ζ*_*t,a*_ = Γ_*a*_ + *B*_*t*_ is the total architecture-dependent translational friction.

Figure 6a,b shows the largest score for each function after optimizing binder allocation with at least one binder and one cleaver. Figure 6c uses three representative states: a polarized explorer and mixed escape specialist share one molecular state, while a polarized exploit specialist occupies a binder-rich state. If state *r* has capacity *U*_*r*_ and population fraction *π*_*r*_, its contribution is *π*_*r*_*U*_*r*_. For prescribed functional demands *d*_*r*_ *>* 0, we chose the allocation

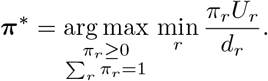

The central point uses equal demands. Varying *d*_explore_*/d*_exploit_ from 1*/*20 to 20 while holding *d*_escape_ fixed produces the sweep in Fig. 6c. The states are projected onto *λ*_bc_ = log_10_[(*ϕ*_*b*_ + 0.003)*/*(*χ*_*C*_ + 0.003)], and each colored area integrates to its population weight. SI Section 6 derives the function scores, weights and projection.

### Statistics

Unless stated otherwise, we first averaged recordings within each biological replicate, giving each recording equal weight. Null intervals came from randomized displacement directions. We summarized simulation– theory agreement by the median absolute difference in log space for conditions in which particles remained mobile. We did not assign a late-time mobility to simulations that remained trapped.

## Data availability

The data supporting the findings of this study are available from the corresponding author upon reasonable request. Data underlying the published figures will be deposited in a public repository before publication.

## Code availability

The simulation and analysis code supporting this study is available to editors and reviewers upon request and will be deposited in a public repository before publication.

## Acknowledgements

We thank Elizabeth May for technical guidance on supported lipid bilayer preparation, and members of the Fletcher laboratory for helpful discussions and feedback throughout this work.

## Author contributions

S.A., G.H. and D.A.F. conceived the study. S.A. developed the theory and simulation framework, carried out the simulations and experiments, analyzed the data and wrote the original manuscript. L.F.O. and D.A.F. contributed to the experimental work. B.V. and G.H. contributed to the theoretical development. B.V., G.H. and D.A.F. contributed to interpreting the results and revising the manuscript. All authors approved the final manuscript.

## References

[1] Einstein, A. Ü ber die von der molekularkinetischen Theorie der Wärme geforderte Bewegung von inruhenden Flüssigkeiten suspendierten Teilchen. Annalen der Physik 322, 549–560 (1905).

[2] von Smoluchowski, M. Zur kinetischen Theorie der Brownschen Molekularbewegung und der Suspensionen. Annalen der Physik 326, 756–780 (1906).

[3] Ramaswamy, S. The mechanics and statistics of active matter. Annual Review of Condensed Matter Physics 1, 323–345 (2010).

[4] Bechinger, C. et al. Active particles in complex and crowded environments. Reviews of Modern Physics 88, 045006 (2016).

[5] de Gennes, P.-G. La percolation: un concept unificateur. La Recherche 7, 919–927 (1976).

[6] Hwang, S.Lee, D.-S. & Kahng, B. Blind and myopic ants in heterogeneous networks. Physical Review E 90, 052814 (2014).

[7] Barbier-Chebbah, A., Bénichou, O. & Voituriez, R. Self-interacting random walks: Aging, exploration, and first-passage times. Physical Review X 12, 011052 (2022).

[8] Solon, J., Streicher, P., Richter, R., Brochard-Wyart, F. & Bassereau, P. Vesicles surfing on a lipid bilayer: Self-induced haptotactic motion. Proceedings of the National Academy of Sciences 103, 12382–12387 (2006).

[9] Lund, K. et al. Molecular robots guided by prescriptive landscapes. Nature 465, 206–210 (2010).

[10] Korosec, C. S. et al. Motility of an autonomous protein-based artificial motor that operates via a burnt-bridge principle. Nature Communications 15, 1511 (2024).

[11] Button, B. et al. A periciliary brush promotes the lung health by separating the mucus layer from airway epithelia. Science 337, 937–941 (2012).

[12] Cohen, M. et al. Influenza A penetrates host mucus by cleaving sialic acids with neuraminidase. Virology Journal 10, 321 (2013).

[13] Vahey, M. D. & Fletcher, D. A. Influenza A virus surface proteins are organized to help penetrate host mucus. eLife 8, e43764 (2019).

[14] Agarwal, S., Veytsman, B., Fletcher, D. A. & Huber, G. Kinetics and optimality of influenza A virus locomotion. Physical Review Letters 133, 248402 (2024).

[15] de Vries, E., Du, W., Guo, H. & de Haan, C. A. M. Influenza A virus hemagglutinin-neuraminidase-receptor balance: Preserving virus motility. Trends in Microbiology 28, 57–67 (2020).

[16] Ziebert, F. & Kulić, I. M. How influenza’s spike motor works. Physical Review Letters 126, 218101 (2021).

[17] Stevens, L., de Buyl, S. & Mognetti, B. M. The sliding motility of the bacilliform virions of Influenza A viruses. Soft Matter 19, 4491–4501 (2023).

[18] Vahey, M. D. & Fletcher, D. A. Low-fidelity assembly of influenza A virus promotes escape from host cells. Cell 176, 281–294.e19 (2019).

[19] Pólya, G. Ü ber eine Aufgabe der Wahrscheinlichkeitsrechnung betreffend die Irrfahrt im Straßennetz. Mathematische Annalen 84, 149–160 (1921).

[20] Berg, H. C. & Purcell, E. M. Physics of chemoreception. Biophysical Journal 20, 193–219 (1977).

[21] Endres, R. G. & Wingreen, N. S. Accuracy of direct gradient sensing by single cells. Proceedings of the National Academy of Sciences 105, 15749–15754 (2008).

[22] Sieben, C., Sezgin, E., Eggeling, C. & Manley, S. Influenza A viruses use multivalent sialic acid clusters for cell binding and receptor activation. PLOS Pathogens 16, e1008656 (2020).

[23] Sakai, T., Nishimura, S. I., Naito, T. & Saito, M. Influenza A virus hemagglutinin and neuraminidase act as novel motile machinery. Scientific Reports 7, 45043 (2017).

[24] Partlow, E. A. et al. Influenza a virus rapidly adapts particle shape to environmental pressures. Nature Microbiology 10, 784–794 (2025).

[25] Peterl, S. et al. Morphology-dependent entry kinetics and spread of influenza a virus. The EMBO Journal 44, 3959–3982 (2025).

[26] Kaler, L. et al. Influenza A virus diffusion through mucus gel networks. Communications Biology 5, 249 (2022).

[27] Iseli, A. N. et al. The neuraminidase activity of influenza A virus determines the strain-specific sensitivity to neutralization by respiratory mucus. Journal of Virology 97, e01271–23 (2023).

[28] Liu, M. et al. Human-type sialic acid receptors contribute to avian influenza A virus binding and entry by hetero-multivalent interactions. Nature Communications 13, 4054 (2022).

[29] Ivanovic, T., Choi, J. L., Whelan, S. P., van Oijen, A. M. & Harrison, S. C. Influenza-virus membrane fusion by cooperative fold-back of stochastically induced hemagglutinin intermediates. eLife 2, e00333 (2013).

[30] Antal, T. & Krapivsky, P. L. “burnt-bridge” mechanism of molecular motor motion. Physical Review E 72, 046104 (2005).

[31] Yehl, K. et al. High-speed DNA-based rolling motors powered by RNase H. Nature Nanotechnology 11, 184–190 (2016).

[32] Saffarian, S., Collier, I. E., Marmer, B. L., Elson, E. L. & Goldberg, G. Interstitial collagenase is a brownian ratchet driven by proteolysis of collagen. Science 306, 108–111 (2004).

[33] Kranz, W. T., Gelimson, A., Zhao, K., Wong, G. C. L. & Golestanian, R. Effective dynamics of microorganisms that interact with their own trail. Physical Review Letters 117, 038101 (2016).

[34] Perl, A. et al. Gradient-driven motion of multivalent ligand molecules along a surface functionalized with multiple receptors. Nature Chemistry 3, 317–322 (2011).

[35] Tweedy, L. et al. Seeing around corners: Cells solve mazes and respond at a distance using attractant breakdown. Science 369, eaay9792 (2020).

[36] LeFebre, R., Landsittel, J. A., Stone, D. E. & Mugler, A. Role of signal degradation in directional chemosensing. Physical Review Letters 133, 138402 (2024).

[37] Kussell, E. & Leibler, S. Phenotypic diversity, population growth, and information in fluctuating environments. Science 309, 2075–2078 (2005).

[38] Beaumont, H. J. E., Gallie, J., Kost, C., Ferguson, G. C. & Rainey, P. B. Experimental evolution of bet hedging. Nature 462, 90–93 (2009).

[39] Landauer, R. Irreversibility and heat generation in the computing process. IBM Journal of Research and Development 5, 183–191 (1961).

[40] Still, S., Sivak, D. A., Bell, A. J. & Crooks, G. E. Thermodynamics of prediction. Physical Review Letters 109, 120604 (2012).

[41] Parrondo, J. M. R., Horowitz, J. M. & Sagawa, T. Thermodynamics of information. Nature Physics 11, 131–139 (2015).

[42] Hoshino, A., Costa-Silva, B., Shen, T.-L., Rodrigues, G. et al. Tumour exosome integrins determine organotropic metastasis. Nature 527, 329–335 (2015).

[43] Shimoda, M. & Khokha, R. Metalloproteinases in extracellular vesicles. Biochimica et Biophysica Acta (BBA) - Molecular Cell Research 1864, 1989–2000 (2017).

[44] Cheng, Q. et al. Selective organ targeting (SORT) nanoparticles for tissue-specific mRNA delivery and CRISPR–Cas gene editing. Nature Nanotechnology 15, 313–320 (2020).

[45] Kaspar, C., Ravoo, B. J., van der Wiel, W. G., Wegner, S. V. & Pernice, W. H. P. The rise of intelligent matter. Nature 594, 345–355 (2021).

[46] Dias, C. S., Trivedi, M., Volpe, G., Araújo, N. A. M. & Volpe, G. Environmental memory boosts group formation of clueless individuals. Nature Communications 14, 7324 (2023).

[47] Olver, F. W. J., Lozier, D. W., Boisvert, R. F. & Clark, C. W. (eds) NIST Handbook of Mathematical Functions (Cambridge University Press, New York, NY, 2010).

[48] Pereira, M. E. A., Kabat, E. A. & Sharon, N. Immunochemical studies on the specificity of soybean agglutinin. Carbohydrate Research 37, 89–102 (1974).

[49] Eisen, M. B., Sabesan, S., Skehel, J. J. & Wiley, D. C. Binding of the influenza A virus to cell-surface receptors: Structures of five hemagglutinin–sialyloligosaccharide complexes determined by X-ray crystallography. Virology 232, 19–31 (1997).

