## Supplementary Information for "Environment-driven active transport of influenza A virus"

Briefly, we consider a rod carrying reversible binders that bind to receptors in the environment and cleavers that irreversibly break them. Binding reads local receptor availability, cleavage writes a persistent change, and ligand placement determines how that change feeds back on particle motion. Polarized particles move mainly along their axes and write narrow trails; mixed particles move mainly across their axes and clear a broader region. The same molecular parameters determine persistence, long-time diffusion, surface exploration, gradient guidance and population transport functions. We use $K_D$ for the unbinding parameter and $K_C$ for cleavage strength; smaller $K_D$ denotes stronger binding.

#### 1 Stochastic particle simulations

The simulations couple discrete molecular reactions to continuous rigid-body motion: the current attachments set the spring forces, motion changes ligand–receptor distances, and the updated geometry sets the next reaction rates.

##### 1.1 Molecular reactions and rigid-body motion

The particle has center $\mathbf{X}$, unit axis $\mathbf{u}$, and ligand positions $\mathbf{r}_i = \mathbf{X} + s_i\mathbf{u}$, with fixed axial coordinates $s_i$. We use $L = 20$ and $N_{\text{tot}} = \text{int}(L) = 20$ axial sites, with their spacing set to one simulation unit. Polarized particles contain contiguous binder and cleaver blocks. In the deterministic mixed layout, cleavers occupy both endpoints and any remaining cleavers are distributed approximately uniformly among the interior sites. Receptor coordinates are held fixed over each trajectory, representing the regime in which redistribution is slower than subsequent rebinding or trail re-encounter. Micron-scale cleaved-sialic-acid trails have been observed in native mucus, and directional bias was retained in simulations that included slow receptor diffusion and boundary replenishment [1]. Receptors occupy one of three states: free, binder-bound or cleaved. A cleaved receptor remains unavailable. Binders attach reversibly, whereas cleavers act directly on nearby free receptors. At ligand–receptor separation $r$, the interaction weight $\omega(r) = e^{-r^2/\alpha^2}$, where $\alpha$ is the interaction reach, gives the reaction rates

$$q_{\text{on}} = \frac{\omega}{1+\omega}, \qquad q_{\text{off}} = \frac{K_D}{1+\omega}, \qquad q_{\text{c}} = K_C\omega.$$

At each fixed geometry, these channels are sampled with the direct stochastic-simulation method [2]. The binding-rate prefactor defines the simulation time unit, so $k_0 \equiv 1$ in these rates. The factor $k_0$ is restored below only when mean-field expressions are written in dimensional time. The ratio $q_{\rm on}/q_{\rm off} = \omega/K_D$ gives the equilibrium binding weight used by the mean-field theory. Possible reaction partners are restricted to receptors within $3\alpha$ in uniform and surface simulations and $2.5\alpha$ in gradient simulations. This local neighborhood is updated as the particle moves, while existing attachments remain tracked outside it.

Each attachment is a zero-rest-length harmonic spring between binder $\mathbf{r}_b$ and receptor $\mathbf{r}_\mu$, with force $\mathbf{F}_b = -k_s(\mathbf{r}_b - \mathbf{r}_\mu)$ and torque $\mathbf{T}_b = s_b\mathbf{u} \times \mathbf{F}_b$. Summing over attachments gives $\mathbf{F}$ and $\mathbf{T}$. The particle has axial and transverse friction coefficients $\Gamma_\parallel$ and $\Gamma_\perp$, and rotational friction $\Gamma_R$. With $\mathbf{P}_\parallel = \mathbf{u}\mathbf{u}^\mathsf{T}$ and $\mathbf{P}_\perp = \mathbf{I} - \mathbf{P}_\parallel$, the translational mobility and diffusion tensors are $\mathbf{M}_t = \mathbf{P}_\parallel/\Gamma_\parallel + \mathbf{P}_\perp/\Gamma_\perp$ and $\mathbf{D}_t = D_\parallel\mathbf{P}_\parallel + D_\perp\mathbf{P}_\perp$. Between reactions, we use overdamped anisotropic rigid-body Brownian dynamics [3, 4],

$$\mathrm{d}\mathbf{X} = \mathbf{M}_t\mathbf{F}\,\mathrm{d}t + \sqrt{2\mathbf{D}_t}\,\mathrm{d}\mathbf{W}_t, \qquad \mathrm{d}\boldsymbol{\theta} = \frac{\mathbf{P}_\perp\mathbf{T}}{\Gamma_R}\,\mathrm{d}t + \sqrt{2D_R^0}\,\mathbf{P}_\perp\,\mathrm{d}\mathbf{W}_R.$$

These equations describe the motion over a short interval. The orientation is updated by a finite rotation, after which $\mathbf{u}$ is normalized and the ligand positions are recalculated.

The friction coefficients reflect the anisotropic drag of an IAV-sized particle. For a 200–300 nm-long, 50–80 nm-wide particle with aspect ratio $\mathcal{A} = L_{\rm phys}/d_{\rm phys}$, the leading drag estimates at local effective viscosity $\eta_{\rm eff}$ are $\zeta_\parallel \simeq 2\pi\eta_{\rm eff}L_{\rm phys}/\log\mathcal{A}$, $\zeta_\perp \simeq 4\pi\eta_{\rm eff}L_{\rm phys}/\log\mathcal{A}$ and $\zeta_R \simeq \pi\eta_{\rm eff}L^3_{\rm phys}/(3\log\mathcal{A})$, up to order-one shape corrections [5, 6]. These estimates give translational drags of order $10^{-7}$–$10^{-4}$ N s m$^{-1}$ and rotational drags of order $10^{-21}$–$10^{-18}$ N m s across the mucus-viscosity range given below. This hierarchy motivates $\Gamma_\perp \simeq 2\Gamma_\parallel$ and comparatively slow rotation. After nondimensionalization, the translational frictions are $\Gamma_\parallel = 0.1$ parallel to the particle axis and $\Gamma_\perp = 0.2$ transverse to it. The rotational friction is $\Gamma_R = 5$.

With $k_s = 1$, the attachment-work scale is $\varepsilon_b = k_s\alpha^2/2 = 0.125$, and the resulting Gaussian weight matches the distance dependence of the reaction rates. Identifying $\varepsilon_b$ with $k_{\rm B}T$ defines the Stokes–Einstein reference diffusivities $D_\parallel^{\rm ref} = \varepsilon_b/\Gamma_\parallel = 1.25$ and $D_\perp^{\rm ref} = \varepsilon_b/\Gamma_\perp = 0.625$, parallel and transverse to the particle axis, respectively. Rotational diffusion has the reference value $D_R^{\rm ref} = \varepsilon_b/\Gamma_R = 0.025$.

Physical units follow from $\mathbf{X}_{\rm phys} = \ell_0\mathbf{X}$ and $t_{\rm phys} = t/k_0$, so that $v_{\rm phys} = \ell_0 k_0 v$, $D_{\rm phys} = \ell_0^2 k_0 D$ and $D_{R,\rm phys} = k_0 D_R$. Around the previously estimated $\ell_0 \simeq 10$ nm HA spacing, we allow $\ell_0 = 10$–$15$ nm. Other estimates span $k_0 \simeq 0.4$–$2\,\mathrm{s}^{-1}$ for association, $k_{\rm off} \simeq 30\,\mathrm{s}^{-1}$ for dissociation, $k_c \simeq 10$–$100\,\mathrm{s}^{-1}$ for cleavage, $S \simeq 0.01$–$1\,k_{\rm B}T\,\mathrm{nm}^{-2}$ for the effective spring stiffness and $\eta_{\rm eff} \simeq 0.1$–$100$ Pa s for the local mucus viscosity [7]. The corresponding friction per ligand-scale segment is $\gamma \simeq 10^{-9}$–$10^{-6}\,\mathrm{kg\,s}^{-1}$, giving $\gamma k_0/S \simeq 10^{-5}$–$10^2$; the dimensionless translational frictions used here lie well inside this range. For $\alpha = 0.5$, identifying $\varepsilon_b$ with $k_{\rm B}T$ gives $S = 2k_{\rm B}T/(\alpha^2\ell_0^2) \simeq 0.15$–$0.33$ pN nm$^{-1}$, inside the estimated stiffness range. These values set a physical scale, while the dimensionless sweeps resolve the wider transport regimes rather than fixing a single biochemical or rheological calibration.

One diffusivity unit then spans $\ell_0^2 k_0 \simeq 4 \times 10^{-5}$–$4.5 \times 10^{-4}\,\mu\mathrm{m}^2\mathrm{s}^{-1}$. The surface-MSD overlay uses $k_0 = 11.1\,\mathrm{s}^{-1}$ only for that comparison. The more robust measure of the cleavage effect is its fold change, which is independent of either conversion: at $K_D = 100$ and $K_C = 10$, $D_{\rm eff}(K_C)/D_{\rm eff}(0) = 12.7$ for polarized particles and 1.8 for mixed particles.

At a one-second lag in human respiratory mucus, muco-inert 219 nm probes diffuse 41-fold more slowly than their Stokes–Einstein value in water, while 549 nm probes are slowed 590-fold and become sterically confined [8]. Their PEG coating suppresses specific mucus adhesion, so this slowdown estimates the steric and rheological background not represented explicitly here. Controlled polymer gels show that diffusivity is governed by the particle-to-mesh-size ratio and falls sharply as the two lengths become comparable [9]. The 200–300 nm long axis of IAV lies within this confinement-sensitive range, although its narrower cross-section and changing orientation make the local hindrance variable.

We treat deterministic response to receptor forces and this unresolved background displacement as independent coarse-grained inputs, setting $D_\parallel = f_T D_\parallel^{\rm ref}$, $D_\perp = f_T D_\perp^{\rm ref}$ and $D_R^0 = f_T D_R^{\rm ref}$. The dimensionless factor $f_T$ scales only the imposed Gaussian displacements and leaves the deterministic friction and molecular rates fixed; it is not a thermodynamic temperature. We use $f_T = 0.01$, retaining nonzero translation and rotation while placing background motion two orders of magnitude below the local Stokes–Einstein reference, as expected near a steric confinement crossover. Since pore geometry and mucus microstructure vary, this value defines a representative constrained regime; the full interval is tested below. Uniform representative trajectories, uniform diffusivity simulations and all gradient simulations use this value, giving $D_\parallel = 0.0125$, $D_\perp = 0.00625$ and $D_R^0 = 2.5 \times 10^{-4}$. Surface-return simulations use $f_T = 0$, and Supplementary Fig. S1 spans $f_T \in \{0, 0.01, 0.1, 1\}$.

With $\tau_{v,a}$ the velocity-correlation time, the run-scale Péclet number $\mathrm{Pe}_{\mathrm{run},a} = v_a\ell_{p,a}/D_{\mathrm{bg},a} = v_a^2\tau_{v,a}/D_{\mathrm{bg},a}$ compares receptor-driven persistence with background diffusion. We use $D_{\mathrm{bg},p} = D_\parallel$ for polarized axial motion and $D_{\mathrm{bg},m} = D_\perp$ for mixed transverse motion. At fixed speed and persistence, $f_T = 0.01$ makes this ratio 100 times its $f_T = 1$ value, placing the simulated runs in the receptor-dominated regime. Supplementary Fig. S1 shows the full response when both quantities also vary.

For fixed attachments and orientation, each translational or small-angle rotational coordinate diffuses in a harmonic potential. The parallel and transverse relaxation rates are $\lambda_\parallel = n_b k_s/\Gamma_\parallel$ and $\lambda_\perp = n_b k_s/\Gamma_\perp$. The small-angle rotational rate is $\lambda_R = k_s \sum_b s_b^2/\Gamma_R$. We use the analytical update of a coordinate $x$ over a time interval [10]:

$$x(t+h) - x_{\mathrm{eq}} = [x(t) - x_{\mathrm{eq}}]e^{-\lambda_x h} + \sqrt{\frac{D_x}{\lambda_x}(1 - e^{-2\lambda_x h})}\, Z,$$

which reduces to free Brownian motion as $\lambda_x \to 0$. After each update, the geometry, forces and torques are recalculated, restoring the coupling between translation and rotation.

##### 1.2 Coupling reactions and Brownian motion

Brownian motion alternates with stochastic reaction events [11]. At the start of each time step, the possible reactions and their total rate $Q = \sum_j q_j$ are recalculated. The sampled waiting time to the next reaction is $\Delta t_{\mathrm{rxn}} = -\log U_1/Q$. Motion is limited to

$$h_{\mathrm{cap}} = \mathrm{clip}_{[h_{\min}, h_{\max}]}\left[\min\left(h_{\max}, \frac{\eta_h}{Q}, h_{\mathrm{mech}}\right)\right], \quad h_{\mathrm{mech}} = \begin{cases} h_{\mathrm{free}}, & n_b = 0, \\ m_b h_0, & n_b > 0, \end{cases}$$

Here $\mathrm{clip}_{[a,b]}$ restricts a value to the interval $[a, b]$, and $h_0$ is the base time step for an attached particle. For a particle without attachments,

$$h_{\mathrm{free}} = \min\left\{\frac{\delta r_{\max}^2}{2(D_\parallel + 2D_\perp)}, \frac{\delta\theta_{\max}^2}{2D_R^0}\right\}.$$

When $Q = 0$, $\eta_h/Q$ and $\Delta t_{\mathrm{rxn}}$ are taken as infinite. The particle advances for the smaller of $\Delta t_{\mathrm{rxn}}$ and $h_{\mathrm{cap}}$. If the motion limit is reached first, the geometry and rates are recalculated before a new waiting time is drawn. If a reaction occurs first, the rates are reevaluated at the event position and the reaction is chosen in proportion to the updated rates. Shorter motion steps limit changes in reaction rates as the particle moves.

The 3D runs with background motion use $h_{\min} = 0.005$, $h_{\max} = 0.75$, $\eta_h = 3$, $\delta r_{\max} = 1.2\alpha$ and $\delta\theta_{\max} = 0.25$ radians. The $f_T = 0$ controls use a simpler update. A reaction is sampled from the current geometry, the particle moves deterministically with its attachment set fixed, and the reaction is then applied. Attachments stretched beyond $5\alpha$ are broken.

Uniform 3D receptors form an unbounded simple-cubic lattice and surface receptors a square lattice in $z = 0$, both with spacing $d_{\mathrm{rec}} = 0.5$ and a receptor at the origin. Every trajectory starts at $\mathbf{X} = (0, 0, 0)$, $\mathbf{u} = \hat{\mathbf{x}}$, with all receptors free and no attachments or cleaved sites. In the physical range above, $L = 20$ represents a 200–300 nm particle and $\alpha = 0.5$ an interaction reach of 5–7.5 nm. Trajectory $j$ uses seed $42 + j$, with the same indexed sequence reused across parameter conditions.

All times are reported in the binding-rate unit defined above. The uniform speed and persistence ensembles contained 64 trajectories per condition and ran to $t = 2 \times 10^4$. The trajectory examples displayed in main-text Fig. 2b ran to $t = 2 \times 10^5$, while the long-time uniform diffusivity ensembles in main-text Fig. 2f,g contained 64 trajectories per condition and ran to $t = 10^6$. Surface ensembles contained 64–256 trajectories per molecular condition and ran until detachment or $t = 10^6$. Gradient ensembles contained 128 trajectories per displayed condition and ran to $t = 5 \times 10^5$ unless a boundary criterion was met earlier; $\mathrm{CI}_{\mathrm{path}}$ was evaluated at $t = 10^5$. The uniform zero-gradient control contained 64 trajectories per condition and was analyzed through $t = 2 \times 10^5$.

The trajectories in main-text Fig. 2b compare $K_C = 0$ with $K_C = 10$ at $K_D = 10$ and 100, all at $f_T = 0.01$. Teal and red trails mark cleaved receptors for polarized and mixed particles. At $K_D = 100$, the mixed particle removes the nearby receptors needed for continued attachment and remains within this receptor-depleted footprint, illustrating the locally depleted state included in the diffusivity theory.

#### 2 Mean-field theory in a uniform receptor region

The mean field follows the same molecular sequence as the simulations. Binding sets the average number and lifetime of receptor attachments, cleavage creates a receptor difference, ligand placement converts that difference into force, and rotation limits the resulting directional run. The cleavage-to-speed calculation extends our earlier 1D theory [7]—here those molecular steps are carried through to 3D diffusion.

The approximation assumes that binding equilibrates within one interaction-range crossing, that individual attachments persist over distances shorter than the scale of receptor variation, and that a directional run lasts longer than one such crossing. These separations allow molecular rates and ligand positions to be converted into transport coefficients.

#### 2.1 Attachment, cleavage and speed

The Gaussian interaction weight samples $\nu = \pi^{3/2}(\alpha/d_{\text{rec}})^3$ receptors per ligand in an intact 3D region. If a fraction $f_{\text{int}}$ of receptors remains, one binder is attached with probability $p_b = f_{\text{int}}\nu/(K_D + f_{\text{int}}\nu)$, giving $\bar{n} = N_b p_b$ attached binders on average. The mean attachment lifetime is $\tau_b = \kappa_b/K_D$, where $\kappa_b = 1 + 2^{-3/2}$. Because an attachment resists displacement throughout its lifetime, it adds the friction

$$B_t = k_s N_b p_b \tau_b = \frac{k_s \kappa_b N_b f_{\text{int}}\nu}{K_D(K_D + f_{\text{int}}\nu)}. \tag{S1}$$

Thus $B_t \sim K_D^{-2}$ at weak binding and $B_t \sim K_D^{-1}$ at strong binding. Subscripts $p$ and $m$ denote the polarized and mixed layouts, with $\Gamma_p = \Gamma_\parallel$ and $\Gamma_m = \Gamma_\perp$. Attachment-induced friction matches the particle friction without attachments at

$$K_{D,a}^{\text{drag}} = \frac{-f_{\text{int}}\nu + \sqrt{(f_{\text{int}}\nu)^2 + 4k_s\kappa_b N_b f_{\text{int}}\nu/\Gamma_a}}{2}.$$

Here $f_b = N_b/(N_b + N_c)$ and $f_c = 1 - f_b$ are the binder and cleaver fractions.

During passage at speed $v_a$, $\Theta_a = \Omega_a/v_a$ is the mean number of cleavage events for a receptor. Independent events leave an intact fraction $e^{-\Theta_a}$. Binders convert the resulting receptor difference into force $F_a$, while slower motion gives cleavers more time to enlarge that difference. Force and speed therefore obey

$$[\Gamma_a + B_t]v_a = F_a(K_D, \Omega_a/v_a). \tag{S2}$$

For a polarized particle, $\Omega_p = k_0 K_C \sqrt{\pi}\alpha N_c$, and the difference between intact receptors ahead and depleted receptors behind gives

$$F_p = 4 f_b f_c \frac{\varepsilon_b}{2\alpha} \log\left(\frac{K_D + \nu}{K_D + \nu e^{-\Theta_p}}\right).$$

For a mixed particle, the two sides have mean cleavage counts $(1 \mp \mu_m)\Theta_m$, where $\mu_m = 4/(3\pi)$ and $\Omega_m = k_0 K_C \sqrt{\pi}\alpha f_c$. Their binding-work difference across $2\mu_m\alpha$ gives

$$F_m = \frac{N_b \varepsilon_b}{2\mu_m \alpha} \log\left[\frac{K_D + \nu e^{-(1-\mu_m)\Theta_m}}{K_D + \nu e^{-(1+\mu_m)\Theta_m}}\right].$$

Both layouts obey $v_a \propto K_C^{1/2}$ at weak cleavage. The polarized receptor contrast then saturates, so its speed approaches a plateau. In the mixed layout, cleavage eventually reaches both sides sampled by the particle, causing the receptor difference and speed to turn over at $\Theta_m^* = (2\mu_m)^{-1}\log[(1+\mu_m)/(1-\mu_m)]$. At weak binding, the polarized affinity dependence contains $\sqrt{r}/(1+r)$, where $r = B_t/\Gamma_\parallel$, and peaks when the two sources of translational friction are comparable.

#### 2.2 Persistence and effective diffusion

Let $s_i$ be the axial binder positions and $I_b = \sum_i (s_i - \bar{s}_b)^2$ their squared spread about the mean. After transverse translation relaxes, a rotation stretches binder $i$ by $(s_i - \bar{s}_b)\,d\theta$. For a symmetric mixed layout,

$$D_R = \frac{D_R^0 \Gamma_R^2 + (k_s\alpha^2/2)B_R}{(\Gamma_R + B_R)^2}, \qquad B_R = k_s p_b \tau_b \sum_i s_i^2.$$

Writing $r_R = B_R/\Gamma_R$, $\tau_0 = (2D_R^0)^{-1}$ and $\Lambda_R = k_s\alpha^2/(2D_R^0\Gamma_R)$ gives $\tau^{\text{run}} = \tau_0(1+r_R)^2/(1+\Lambda_R r_R)$. At weak binding, $r_R \simeq k_s\kappa_b\nu\sum_i s_i^2/(\Gamma_R K_D^2)$, and hence

$$\tau^{\text{run}} \simeq \tau_0\left[1 - (\Lambda_R - 2)\frac{k_s\kappa_b\nu\sum_i s_i^2}{\Gamma_R K_D^2}\right].$$

For the chosen background-motion parameters, $\Lambda_R > 2$, so this branch falls from the weak-binding rotational limit as affinity strengthens and reaches its minimum at $r_R^* = 1 - 2/\Lambda_R$, close to one. In the strong-binding limit,

$$D_{R,a} \simeq \frac{\alpha^2 K_D}{2\kappa_b I_{b,a}}, \qquad \tau_a^{\mathrm{run}} \simeq \frac{\kappa_b I_{b,a}}{\alpha^2 K_D}. \tag{S3}$$

The strong-binding persistence is therefore the attachment lifetime multiplied by the binder lever-arm factor $I_b/\alpha^2$. The polarized calculation retains translation–rotation coupling but has the same strong-binding dependence on $I_b$. The polarized architecture has a vectorial orientation, so its lifetime in main-text Fig. 2d is $(2D_{R,p})^{-1}$. The mixed architecture is apolar, so the plotted lifetime is $(6D_{R,m})^{-1} = \tau_m^{\mathrm{run}}/3$.

We write $D_{\mathrm{bu},a}$ for the cleavage-free diffusion generated by background fluctuations and reversible binding–unbinding events. For a polarized particle, $\tau_{v,p} = \tau_p^{\mathrm{run}}$. A mixed velocity can also decorrelate by directed or diffusive crossing of the body-wide receptor region, so

$$(\tau_{v,m})^{-1} = (\tau_m^{\mathrm{run}})^{-1} + \frac{v_{\mathrm{coh},m}}{L+2\alpha} + \frac{8D_\perp}{(L+2\alpha)^2}.$$

Here $P_{\mathrm{att}} = 1-(1-p_b)^{N_b}$ is the probability of at least one attachment, and $v_{\mathrm{coh},m}^2 = P_{\mathrm{att}} v_m^2$ is the corresponding attachment-weighted squared speed. Let $v_{\mathrm{rms},p}^2$ denote the second moment of polarized speed, and let $\sigma_{v,m}^2$ denote the mixed speed variance. Persistent motion then adds

$$D_{\mathrm{act},p} = \frac{v_{\mathrm{rms},p}^2 \tau_{v,p}}{3}, \qquad D_{\mathrm{act},m} = \frac{v_{\mathrm{coh},m}^2 \tau_{v,m} + \sigma_{v,m}^2 \tau_m^{\mathrm{run}}}{3}. \tag{S4}$$

Fluctuations in the number and placement of attached binders give $\sigma_{v,m}^2 = 4p_b(1-p_b)c^3 v_{\mathrm{coh},m}^2$, with $c = 2\min(f_b, f_c)$. Here $c^3$ approximates the probability that the finite ligand pattern contains both ligand types along all three spatial directions. The polarized correction follows from the same average over the discrete ligand pattern.

These expressions select an intermediate affinity: weak binding transmits little receptor-generated force, whereas strong binding creates large attachment-induced friction. For polarized particles at fixed $K_C$, $D_{\mathrm{act},p} \sim K_D \log^2(\nu/K_D)$ as $K_D \to 0$, so the active contribution tends to zero at strong binding. When persistence varies slowly, the leading persistent-motion optimum lies near $K_D^{\mathrm{drag}}$. Its shift obeys $2\,\partial \log v/\partial \log K_D + \partial \log \tau/\partial \log K_D = 0$. The full optimum also includes the variation of $D_{\mathrm{bu}}$.

#### 2.3 Local receptor depletion and the diffusivity optimum

For either architecture, we define the cleavage-induced diffusivity shift as $\Delta D_{\mathrm{eff},a} = D_{\mathrm{eff},a}(K_C) - D_{\mathrm{eff},a}(0)$. After reversal, a mixed particle can enter the receptor-poor region produced during its preceding run. This region has width $w_m = L + 2\alpha$. Averaging cleavage over its diffusive crossing time gives

$$E_m = \frac{k_0 K_C \nu (6\alpha f_c^3) w_m}{8 D_{\mathrm{bu},m}}, \qquad f_{\mathrm{dep}} = e^{-E_m}.$$

The probability of complete unbinding upon entry, conditional on prior attachment, is

$$\psi_m = \frac{[K_D/(K_D + f_{\mathrm{dep}}\nu)]^{N_b} - [K_D/(K_D+\nu)]^{N_b}}{1 - [K_D/(K_D+\nu)]^{N_b}}.$$

A run of mean length $v_{\mathrm{coh},m}\tau_{v,m}$ crosses this region with probability $P_{\mathrm{cross}} = e^{-w_m/(v_{\mathrm{coh},m}\tau_{v,m})}$. The entry rate, exit rate and their ratio are

$$k_{\mathrm{in}} = \frac{P_{\mathrm{cross}}\psi_m}{\tau_{v,m}}, \qquad k_{\mathrm{out}} = \frac{8D_\perp}{w_m^2}, \qquad \Lambda_{\mathrm{dep}} = \frac{k_{\mathrm{in}}}{k_{\mathrm{out}}}.$$

Here $k_{\mathrm{in}}$ is the rate of losing all attachments in the locally depleted region, while $k_{\mathrm{out}}$ is the rate of returning to intact receptors. The crossing time used to calculate receptor depletion is set by the orientationally averaged $D_{\mathrm{bu},m}$, whereas departure from the body-wide depleted region is transverse and therefore set by $D_\perp$. The ratio $\Lambda_{\mathrm{dep}}$ measures the relative residence in the locally depleted state. The corresponding mobile fraction is $f_{\mathrm{mob},m} = 1/(1+\Lambda_{\mathrm{dep}})$. The mixed diffusivity shift relative to the matched cleavage-free particle is

$$\Delta D_{\mathrm{eff},m} = f_{\mathrm{mob},m} D_{\mathrm{act},m} - (1 - f_{\mathrm{mob},m}) D_{\mathrm{bu},m}. \tag{S5}$$

It crosses zero at $\Lambda_{\rm dep} = D_{{\rm act},m}/D_{{\rm bu},m}$. At weak cleavage, $D_{{\rm act},a} \propto K_C$. Polarized motion then approaches a plateau as receptor contrast saturates. Mixed motion has an interior optimum because the transverse contrast turns over while the locally depleted fraction continues to grow. Composition creates a related trade-off: binders transmit the receptor difference, while cleavers create it, and either function is lost when one ligand type becomes too sparse.

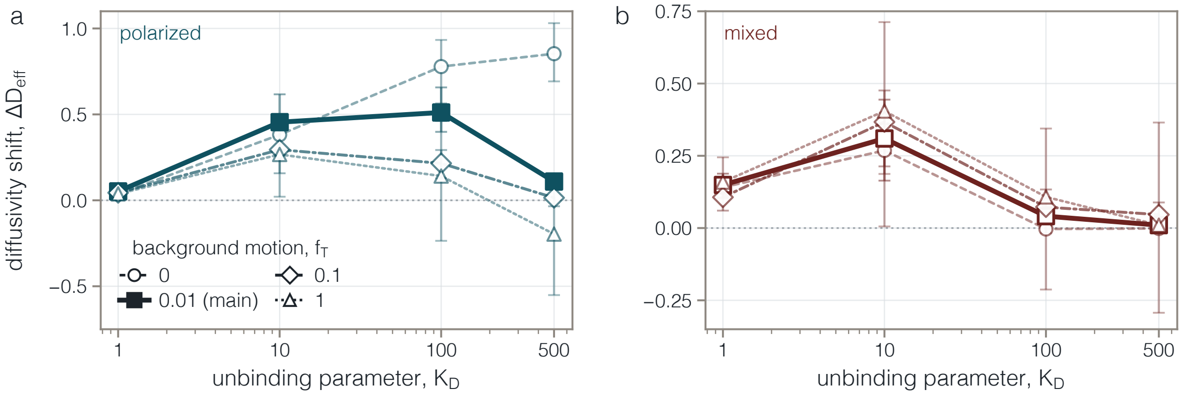


**Supplementary Figure S1 Receptor-driven diffusivity enhancement persists across background-motion strengths. a,b**, Cleavage-induced diffusivity shift $\Delta D_{\rm eff} = D_{\rm eff}(K_C = 10) - D_{\rm eff}(K_C = 0)$ versus $K_D$ for polarized (**a**) and mixed (**b**) particles at background-motion factors $f_T = 0, 0.01, 0.1, 1$. Smaller $K_D$ corresponds to stronger attachment; the $f_T = 0.01$ series used in the main simulations is emphasized. The factor $f_T$ scales the imposed translational and rotational Gaussian motion from the zero-background limit ($f_T = 0$) to the local Stokes–Einstein reference ($f_T = 1$). Points are differences between ensemble mean-squared-displacement slopes, and lines guide the eye. Error bars show 95% bootstrap intervals obtained by resampling complete trajectory ensembles. For each $K_D$, $n = 128$ trajectories per $K_C$ condition at $f_T = 0$, and $n = 64$ at each nonzero $f_T$. The $f_T = 0$ runs extend to $t = 10^6$, and the nonzero-$f_T$ runs to $t = 2 \times 10^5$.

The influence of background motion is set by multivalent attachment. In an intact 3D receptor region, $p_b = \nu/(K_D + \nu)$, so the average number of attached binders is $\bar{n} = N_b\nu/(K_D + \nu)$, and the probability of at least one attachment is $P_{\rm att} = 1 - (1 - p_b)^{N_b}$. The crossover $\bar{n} \simeq 1$ occurs at $K_D^{\rm att} \simeq (N_b - 1)\nu$. For $N_b = 10$ and $\alpha = d_{\rm rec} = 0.5$, $\nu = \pi^{3/2} \simeq 5.57$, giving $K_D^{\rm att} \simeq 50$; $P_{\rm att}$ is 0.99 at $K_D = 10$, but falls to 0.42 at $K_D = 100$. Below this crossover, attachment springs confine the imposed stochastic displacements, and the cleavage-induced diffusivity shifts at the sampled $K_D = 1$ and 10 show no resolved dependence on $f_T$ within their uncertainty intervals (Supplementary Fig. S1). At weaker binding, background rotation can interrupt polarized runs, while transverse background motion can help mixed particles leave locally depleted regions. The choice of $f_T$ is therefore subdominant in the multivalently bound regime and becomes a physical control parameter once complete detachment is common.

In 3D, returns to a bounded region do not accumulate with probability one, so the most recently depleted region gives the leading correction. On a surface, encounters with trails written many runs earlier accumulate, as derived next.

### 3 Surface returns and exploration range

A particle wandering on a 2D surface repeatedly returns to neighborhoods visited earlier, a classical property of planar random walks [12–14]. Cleavage turns each visit into a persistent receptor trail whose accumulated effect is set by the persistence length $\ell_p$ of one directional run and its effective cleaved-trail width $w$. Unless stated otherwise, the symbols refer to the architecture being evaluated.

The surface simulations use $K_D = 10$ and $f_T = 0$ to isolate receptor-mediated motion and repeated encounters with the written trail. At this affinity, the 3D $f_T$ sweep shows that the positive diffusivity shift generated by cleavage changes little with $f_T$ (Supplementary Fig. S1). Detachment is the terminal loss of receptor contact: after the last attachment breaks, the trajectory ends only when no intact receptor remains accessible for rebinding. The center-of-mass displacement at that point defines the terminal exploration range. Trajectories that remained attached at $t = 10^6$ contributed their displacement at that time to the reported mean.

##### 3.1 Single-run transport and cleaved-trail width

On a plane, one ligand samples $\nu = \pi(\alpha/d_{\text{rec}})^2$ receptors. We use this planar receptor number with the attachment lifetime $\tau_b = \kappa_b/K_D$ defined in the uniform theory. The resulting binding–unbinding diffusion is

$$D_{\text{bu}} = \frac{N_b p_b \sigma_F^2 \tau_b}{2}\left[(\Gamma_\parallel + B_t)^{-2} + (\Gamma_\perp + B_t)^{-2}\right], \qquad \sigma_F^2 = \frac{k_s^2\alpha^2}{2}.$$

The surface speed follows the same molecular force balance as the uniform theory with this planar value of $\nu$. Planar rotation gives $\tau_p = D_R^{-1}$, while a mixed particle also exchanges its transverse direction as it moves through the receptor landscape, giving $\tau_m = [D_R + \sqrt{2/\pi}\, v_{\text{rms}}/\alpha]^{-1}$. For either layout,

$$D = D_{\text{bu}} + \frac{v_{\text{rms}}^2\tau}{2}, \qquad \ell_p = \sqrt{2D\tau}.$$

One cleaver acts over Gaussian area $\pi\alpha^2$. Dividing the total area changed per unit time among the $N_b$ statistically equivalent, binder-coupled portions of the path gives $\dot{A}_c = \pi\alpha^2 k_0 K_C (N_c/N_b)$. The available geometric width is $w_{\text{geom}} = 2\alpha$ for a polarized particle and $w_{\text{geom}} = L_c + 2\alpha$ for a mixed particle, where $L_c$ is the axial span of its cleavers. Distributing this area over the footprint of one run gives

$$q = \frac{\dot{A}_c\tau}{\ell_p w_{\text{geom}}}, \qquad w = w_{\text{geom}}(1 - e^{-q}). \tag{S6}$$

Thus $w \simeq \dot{A}_c\tau/\ell_p$ at weak cleavage and $w \simeq w_{\text{geom}}$ once the footprint is filled.

##### 3.2 Repeated returns and exploration range

After $j$ further runs, the probability of a detachment-inducing encounter with one old trail is $w/[2\pi\ell_p(j+1)]$. Summing this probability over all trail ages and approximating the discrete sums by integrals gives $H(n) = (w/2\pi\ell_p)[(n+1)\log(n+1) - n]$. The fraction still attached after $n$ runs is $e^{-H(n)}$, while its mean radial distance is $\sqrt{\pi D\tau n}$. Setting $H(n_*) \simeq 1$ for $n_* \gg 1$ gives the characteristic exploration range

$$R_{\text{MF}} = \frac{\pi\ell_p}{\sqrt{(w/\ell_p)W_0(2\pi\ell_p/w)}}. \tag{S7}$$

Here $W_0$ is the principal branch of the Lambert $W$ function. The complete path history therefore enters through one dimensionless control parameter, $w/\ell_p$. For a narrow trail,

$$R_{\text{MF}} \simeq \pi\sqrt{\frac{\ell_p^3}{w\log(2\pi\ell_p/w)}}.$$

Persistence helps twice by extending each run and spreading old trails over a larger area, which produces the $\ell_p^{3/2}/\sqrt{w}$ dependence.

At weak cleavage, this control parameter reduces to molecular inputs,

$$\frac{w}{\ell_p} \simeq \frac{\pi\alpha^2 k_0 K_C}{2D}\frac{N_c}{N_b}.$$

The range then scales as $K_C^{-1/2}$, apart from the logarithm, when $D$ and $\tau$ take their leading $K_C \to 0$ values. At strong cleavage, one inserts $w = 2\alpha$ for the polarized layout or $w = L_c + 2\alpha$ for the mixed layout into equation S7. Particle length therefore enters when the mixed footprint fills.

For the comparison in main-text Fig. 3d, molecular rates and ligand geometry determine $D$, $\tau$, $\ell_p$ and $w$. Equation S7 then gives the predicted range, which is compared with the simulation mean and the interquartile range across trajectories without fitting the range data.

### 4 Transport along a receptor-density gradient

The 3D gradient simulations add the simplest spatial heterogeneity while retaining the uniform reaction, friction and $f_T = 0.01$ background-motion parameters. Receptor spacing remains $d_{\text{rec}}$ along two axes. Along the third, successive plane spacings are multiplied by $g$ on one side of the origin and divided by $g$

on the other. For $g > 1$, the side with successively smaller spacings is denser. Spacings were restricted to $0.1 \le d \le 5$. Trajectories stopped when the minimum receptor spacing within their ligand-accessible interval fell below 0.12 on the dense side, while a sparse-side exit required a plane crossing followed by a 1,000-unit dwell. These simulation limits are distinct from the absorbing boundaries used below to define path and arrival observables.

We orient $z$ toward denser receptors and define $s_\rho = d/d_{\rm rec}$. Before either spacing limit is reached, the local spacing is well described by $s_\rho(z) = 1 - \beta z$, with $\beta = (g-1)/d_{\rm rec} \simeq \log(g)/d_{\rm rec}$, and the relative density is $\rho(z) = 1/s_\rho(z)$. When $\beta\alpha \ll 1$ and $\beta L \ll 1$, a ligand samples $\nu_\rho = \nu/s_\rho$ receptors without changing $K_D$ or $K_C$. Density at the particle center sets local speed and diffusion, and the difference between the two ends sets the preferred direction.

#### 4.1 Local drift and directional bias

The Gaussian interaction weight sets the force–displacement work scale $\varepsilon_b = k_s\langle x^2\rangle = k_s\alpha^2/2$. A fractional density change alters the number of accessible receptors by the same fraction. Integrating the resulting attachment work from the sparse to the dense end of the particle gives the density-driven force. Balancing that force against particle and attachment-induced friction gives the small-gradient drift $u_{\rm bind} = \beta U_{\rm bind}$, where

$$U_{\rm bind}(s_\rho) = \frac{\varepsilon_b N_b \nu}{\Gamma_\parallel K_D s_\rho^2 + \Gamma_\parallel \nu s_\rho + k_s\kappa_b N_b \nu s_\rho/K_D}. \tag{S8}$$

The drift scales as $K_D^{-1}$ when attachments are sparse and as $K_D$ when attachment-induced friction dominates. At fixed $s_\rho$, it peaks at $K_D^*(s_\rho) = \sqrt{k_s\kappa_b N_b\nu/(\Gamma_\parallel s_\rho)}$. The plotted curves average force times mobility over the binomial distribution of attached binders.

For polarized particles, the uniform speed generated by cleavage, $v_{\rm cleave}$, also acquires a signed projection. At each position, that speed is evaluated with receptor count $\nu_\rho = \nu/s_\rho$ and the original molecular $K_D$ and $K_C$. Writing $p_{b,+}$ and $p_{b,-}$ for the attachment probabilities at the dense and sparse ends, their normalized difference is

$$\eta = \frac{p_{b,+} - p_{b,-}}{p_{b,+} + p_{b,-}} \simeq \frac{\beta L K_D}{2(K_D s_\rho + \nu)}.$$

It approaches $\beta L/(2s_\rho)$ at weak binding and tends linearly to zero with $K_D$ at strong binding as attachment probabilities at both ends approach one. Mixed symmetry gives no leading signed projection from cleavage. The uniform theory gives the mixed mobile fraction $f_{{\rm mob},m} = (1+\Lambda_{\rm dep})^{-1}$, so

$$\frac{u_p}{\beta} \simeq U_{\rm bind} + v_{\rm cleave}\frac{L K_D}{2(K_D s_\rho + \nu)}, \qquad \frac{u_m}{\beta} \simeq f_{{\rm mob},m} U_{\rm bind}. \tag{S9}$$

At $K_C = 0$, $v_{\rm cleave} = 0$ and $f_{{\rm mob},m} = 1$, so the two layouts share the same leading drift. We write the total local drift as $u_a = \beta U_a$.

Each trajectory is recorded at 256 equally spaced times. The corresponding mean-field interval $\Delta$ is the expected time before reaching a boundary divided by 255. Let $D_{{\rm path},a}$ be the local displacement diffusion over this interval, including baseline diffusion and the speed variance retained over $\Delta$. When directed progress is small compared with the 3D step size, averaging this ratio over trajectories gives

$${\rm CI}_{{\rm path},a} \simeq \frac{\sqrt{\pi}}{4} u_a \sqrt{\frac{\Delta}{D_{{\rm path},a}}}. \tag{S10}$$

${\rm CI}_{\rm path}$ is signed progress along the gradient divided by path length accumulated in all three directions. The term “chemotactic” denotes geometric alignment and does not imply internal signaling. For binding-dominated motion with gently varying $D_{\rm path}$,

$$K_{D,{\rm CI}} \simeq \sqrt{\frac{k_s\kappa_b N_b \nu}{\Gamma_\parallel}}.$$

The exact shift obeys $2\,\partial_{\log K_D} \log u_a = \partial_{\log K_D} \log D_{{\rm path},a}$. For the plotted prediction, we use the exact mean length of an isotropic 3D Gaussian step and calculate the mean signed displacement and path length until the particle reaches either boundary. Simulations average $\Delta z_i/\mathcal{L}_i$ across trajectories, while the mean field divides the calculated mean progress by the mean path length.

#### 4.2 Arrival at the dense boundary

Over an observation time $T_{\text{arr}}$, let $q_a(z,t)$ be the probability that a particle starting at $z$ reaches the dense boundary by time $t$ before reaching the sparse boundary. It satisfies $\partial_t q_a = u_a \partial_z q_a + D_{\text{arr},a} \partial_z^2 q_a$, with absorbing boundary values one at the dense side and zero at the sparse side. The normalized dense-side arrival rate is

$$J_{\text{arr},a} = T_{\text{arr}} \int_0^{T_{\text{arr}}} \frac{\partial_t q_a(z_0,t)}{t}\,\mathrm{d}t.$$

The prefactor $T_{\text{arr}}$ makes this inverse-time-weighted rate dimensionless. Arrival samples the long-time local diffusivity over a complete crossing. Let $L_g$ be the initial distance to the dense boundary. When the coefficients are nearly constant, the sparse boundary is remote on the first-arrival time scale, and the observation interval exceeds the typical travel time,

$$J_{\text{arr},a} \simeq T_{\text{arr}} \left( \frac{\beta U_a}{L_g} + \frac{2D_{\text{arr},a}}{L_g^2} \right). \tag{S11}$$

The first term rewards directed completion, and the second rewards successful diffusive exploration. The second term keeps arrival finite at zero density contrast, where equation S10 gives $\text{CI}_{\text{path}} = 0$.

The binding-drift travel time gives a closed affinity optimum. If the dense boundary has relative spacing $s_{\rho,a} < 1$, integration of equation S8 yields a sparse-attachment cost proportional to $K_D(1 - s_{\rho,a}^3)/3$ and an attachment-friction cost proportional to $(1 - s_{\rho,a}^2)/(2K_D)$. Their balance gives

$$K_{D,\text{arr}}^{(0)} = \sqrt{\frac{3k_s\kappa_b N_b \nu(1+s_{\rho,a})}{2\Gamma_\parallel(1+s_{\rho,a}+s_{\rho,a}^2)}} = K_{D,\text{CI}} \sqrt{\frac{3(1+s_{\rho,a})}{2(1+s_{\rho,a}+s_{\rho,a}^2)}}. \tag{S12}$$

Since $s_{\rho,a} < 1$, this drift-only traversal optimum lies at weaker attachment than the local CI estimate. It does not determine the ordering of the maxima in main-text Fig. 4g,h, because the full arrival rate also contains the positive diffusivity term in equation S11. In the full mean-field calculation, diffusive spreading adds to the path length that penalizes $\text{CI}_{\text{path}}$, but helps the particle reach the denser boundary. The CI maximum consequently occurs at smaller $\bar{n}/K_D$ than the arrival maximum for the plotted parameters. These curves use the same molecular inputs to evaluate polarized drift, the mixed mobile fraction, spatially varying diffusion, finite observation time and both absorbing boundaries.

### 5 Experimental gradient sensing on measured receptor landscapes

We measured the local response predicted by the gradient theory on supported lipid bilayers displaying sialylated CEACAM5 crosslinked with soybean agglutinin (SBA), which created heterogeneous receptor landscapes. Virions were tracked in the 561-nm channel, the 647-nm CEACAM5 intensity $I(\mathbf{r})$ mapped the receptor landscape, and the 488-nm SBA channel provided an independent image of the same surface structure. We denote untreated virions by NAI− and virions treated with 100 $\mu$M oseltamivir by NAI+. The label noSBA denotes uncrosslinked control bilayers. Viral labeling, SLB preparation and all subsequent assay steps were repeated independently in three biological replicates. Measurements were averaged first within tracks and recordings, then compared across replicates. The SBA data comprised 24 NAI− recordings (12,412 tracks) and 18 NAI+ recordings (13,722 tracks). The noSBA data comprised 16 NAI− recordings (3,000 tracks) and 11 NAI+ recordings (8,260 tracks).

#### 5.1 Particle-scale receptor contrast and receptor-landscape stability

The illumination-corrected 647-nm image was Gaussian-smoothed with $\sigma_g = 0.25$ $\mu$m. At each displacement origin $\mathbf{r}_i$, intensity was sampled around a ring of radius $r_g = 0.5$ $\mu$m:

$$\bar{I}_i = \langle I(\mathbf{r}_i + r_g \hat{\mathbf{e}}_\varphi) \rangle_\varphi, \qquad \mathbf{h}_i = \langle I(\mathbf{r}_i + r_g \hat{\mathbf{e}}_\varphi)\hat{\mathbf{e}}_\varphi \rangle_\varphi, \qquad C_i = \frac{4|\mathbf{h}_i|}{\bar{I}_i}.$$

The vector $\mathbf{h}_i$ points toward locally denser receptors. For a linear receptor gradient, $C_i$ is the fractional front-to-back intensity change across the $2r_g = 1.0$ $\mu$m ring diameter. Figures report $100C_i$ in percent. Non-overlapping particle-scale displacements were constructed by following each trajectory until its displacement first reached or exceeded 0.5 $\mu$m:

$$c_i = \frac{\Delta\mathbf{r}_i \cdot \mathbf{h}_i}{|\Delta\mathbf{r}_i|\,|\mathbf{h}_i|} = \cos\theta_i, \qquad \text{CI}_{\text{step}} = \langle c_i \rangle.$$

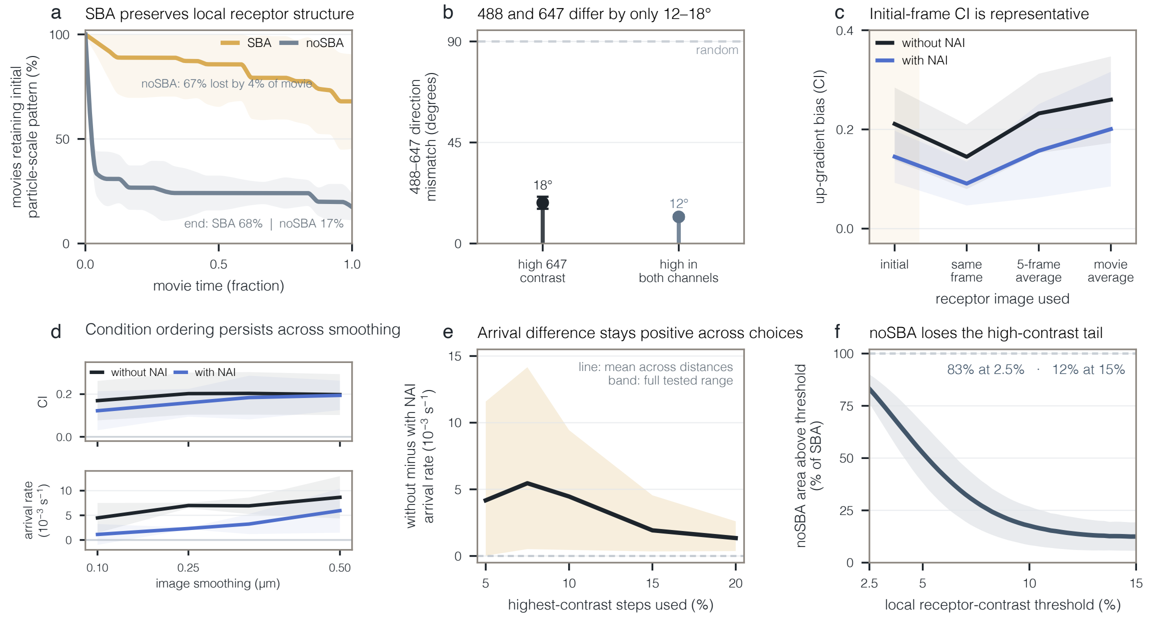


**Supplementary Figure S2 SBA stabilizes local receptor structure, and gradient guidance is robust.** **a**, Fraction of recordings retaining the initial particle-scale 647-nm receptor pattern after scalar brightness correction. SBA-crosslinked receptor landscapes remain more stable than noSBA controls. **b**, Direction mismatch between the independent 488-nm SBA and 647-nm CEACAM5 images at high-contrast locations. The dashed line marks the 90° expectation for unrelated directions. **c**, Up-gradient bias $\mathrm{CI}_{\mathrm{step}}$ calculated from alternative receptor images; the initial image preserves the treatment ordering. **d**, $\mathrm{CI}_{\mathrm{step}}$ and net arrival rate $J_{\mathrm{net}}$ for NAI− and NAI+ across receptor-image smoothing scales. **e**, Treatment difference in $J_{\mathrm{net}}$ across the fraction of highest-contrast steps used. The line is the mean across five arrival distances, and the band spans their full range. **f**, High-contrast area in noSBA controls relative to SBA-crosslinked samples as the receptor-contrast threshold increases. The dashed line marks equal areas. Curves and bars show means across three biological replicates; light shading shows s.d. where displayed. Multiple recordings from the same replicate were averaged before comparison.

We fixed the strongest-10% threshold by pooling receptor contrasts from all accepted steps across treatments and biological replicates. The resulting $100C_i = 9.2\%$ cutoff was applied unchanged to every condition. For the arrival analysis, consecutive selected displacements formed a sequence whose progress along the changing local gradient direction was accumulated as $x_n^{\mathrm{cue}} = \sum_{k=1}^{n} \Delta\mathbf{r}_k \cdot \widehat{\mathbf{h}}_k$. First crossing of $+\ell_{\mathrm{arr}}$ or $-\ell_{\mathrm{arr}}$, with $\ell_{\mathrm{arr}} = 0.75\ \mu$m, contributed $+1/t_{\mathrm{hit}}$ or $-1/t_{\mathrm{hit}}$, while sequences reaching neither boundary contributed zero. Their mean, $J_{\mathrm{net}}$, is the net first-arrival rate. The experimental $\mathrm{CI}_{\mathrm{step}}$ and $J_{\mathrm{net}}$ use the same positive, dense-side direction as the theoretical $\mathrm{CI}_{\mathrm{path}}$ and $J_{\mathrm{arr}}$. Their normalizations differ because $\mathrm{CI}_{\mathrm{path}}$ uses the full path length and $J_{\mathrm{arr}}$ counts dense-side arrivals without subtracting sparse-side arrivals.

The initial receptor image remained a suitable reference because SBA crosslinking preserved the particle-scale pattern through each recording. After scalar brightness correction, the fractions retaining the initial pattern at the end of the recording were $68 \pm 21\%$ with SBA and $17 \pm 5\%$ without SBA (mean $\pm$ s.d. across three biological replicates; Supplementary Fig. S2a). This separation persisted across nearby retention criteria. The independent 488-nm SBA and 647-nm CEACAM5 images also recovered nearly the same local direction, with mismatches of $18 \pm 3°$ at high-647-contrast locations and $12 \pm 1°$ where both channels had high contrast (Supplementary Fig. S2b).

The guidance result was likewise insensitive to how the receptor image was processed. Alternative initial, displacement-matched, five-frame-average and recording-average images preserved the ordering of $\mathrm{CI}_{\mathrm{step}}$, so we used the initial image as an unbleached fixed reference (Supplementary Fig. S2c). Both $\mathrm{CI}_{\mathrm{step}}$ and $J_{\mathrm{net}}$ retained their treatment ordering across receptor-image smoothing scales, and the treatment difference in $J_{\mathrm{net}}$ remained positive across the tested highest-contrast fractions and arrival distances (Supplementary Fig. S2d,e).

The noSBA controls provided a complementary spatial check. On a uniform image grid sampled at the same particle scale, the noSBA controls lacked the high-contrast tail created by crosslinking: their area above threshold fell from 83% of the corresponding SBA area at 2.5% contrast to 12% at 15% contrast (Supplementary Fig. S2f).

#### 5.2 Tracking and motility controls

Virions were detected with a background-subtracted Hessian spot detector using a minimum signal-to-noise ratio of 2.0. Tracks used a 1.04 $\mu$m maximum link displacement, three-frame memory and at least 20 detections. Four fast noSBA NAI$-$ recordings used a 1.30 $\mu$m link displacement. Across conditions, 89%–92% of links joined detections in adjacent frames, and the 99th-percentile displacement occupied 51%–71% of the allowed link distance (Supplementary Fig. S3a). Lowering the signal-to-noise threshold from 2.0 to 1.0 changed the yield of tracks with at least 50 detections by less than 1%. Increasing the maximum link displacement from 8 to 14 pixels (1.04–1.82 $\mu$m) did not materially increase long-track yield while preserving adjacent-frame continuity (Supplementary Fig. S3b,c). In noSBA controls, fewer than 0.6% of adjacent-frame displacements were within 20% of the link limit (Supplementary Fig. S3d). The selected thresholds therefore preserved track yield without truncating the displacements of rapidly moving particles.

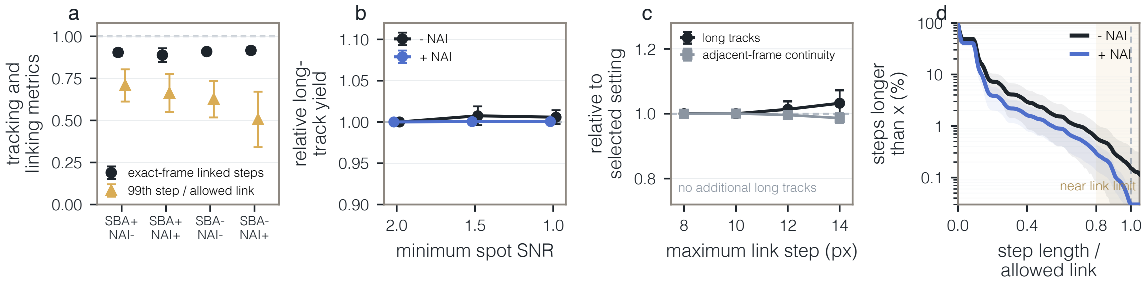


**Supplementary Figure S3 Tracking remains stable across detection and linking choices.** **a**, Fraction of adjacent-frame links and 99th-percentile displacement relative to the allowed link distance for each condition. **b**, Long-track yield across signal-to-noise thresholds in noSBA recordings, normalized to the selected setting. **c**, Long-track yield and adjacent-frame continuity across maximum link displacements in noSBA NAI$-$ recordings. **d**, Adjacent-frame displacement distributions in noSBA controls, normalized by the allowed link distance. Shading marks displacements within 20% of the limit. Points and error bars in **a**–**c**, and lines and shading in **d**, show mean $\pm$ s.d. across three biological replicates. Black denotes NAI$-$ and indigo blue denotes NAI+ where conditions are separated.

#### 5.3 NA inhibition separates alignment from arrival

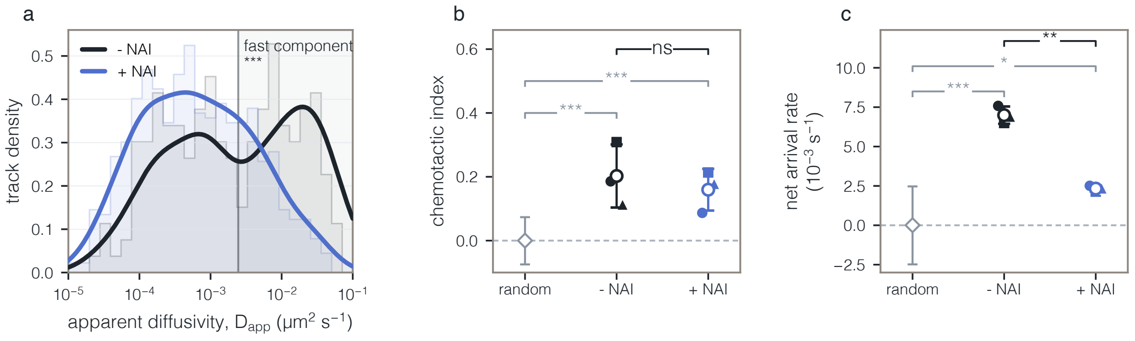


**Supplementary Figure S4 NA inhibition suppresses transport more strongly than local alignment.** **a**, noSBA $D_{\mathrm{app}}$ distributions for NAI$-$ and NAI+. Solid curves average the three biological-replicate-specific density estimates equally, faint histograms show pooled tracks, and shading marks the fast component defined from the untreated distribution. Asterisks show the permutation test obtained by shuffling treatment labels among recordings within each replicate. **b**, $\mathrm{CI}_{\mathrm{step}}$ among the shared strongest 10% of receptor contrasts. Both conditions lie above the direction-randomized null, whereas their difference is unresolved. **c**, $J_{\mathrm{net}}$ in the same window with $\ell_{\mathrm{arr}} = 0.75$ $\mu$m, showing the reduction after NA inhibition. Filled symbols in **b,c** are biological replicates; open circles and error bars show mean $\pm$ s.d. Gray symbols and intervals show direction randomization averaged within recordings and then across biological replicates. Gray brackets compare each condition with its randomized null, and treatment brackets compare paired biological-replicate means. $*P < 0.05$, $**P < 0.01$, $***P < 0.001$; ns, $P \geq 0.05$. Black denotes NAI$-$ and indigo blue denotes NAI+.

Apparent diffusivity $D_{\mathrm{app}}$ was measured on noSBA bilayers by fitting the time-averaged mean-squared displacement over lags 1–10 to $\mathrm{MSD}(\tau) = 4D_{\mathrm{app}}\tau + b$. The analyzed distributions contained 180 NAI$-$ tracks and 606 NAI+ tracks. A two-component Gaussian mixture was fitted to pooled untreated $\log_{10} D_{\mathrm{app}}$, then held fixed when assigning both conditions. The mean of the three biological-replicate-specific probabilities of belonging to the fast component decreased from 0.54 for NAI− to 0.28 for NAI+, with the same direction in all three replicates (Supplementary Fig. S4a). A paired two-sided $t$-test across the three biological-replicate means gave $P = 0.046$, and a permutation test that shuffled treatment labels among recordings within each replicate gave $P < 10^{-4}$.

At the shared cutoff selecting the strongest 10% of receptor contrasts, $\mathrm{CI}_{\mathrm{step}}$ was $0.20 \pm 0.10$ for NAI− and $0.16 \pm 0.07$ for NAI+. Both exceeded the direction-randomized null, but their paired difference was not resolved ($P = 0.51$; Supplementary Fig. S4b). Over the same contrast window, $J_{\mathrm{net}}$ decreased from $(7.0 \pm 0.6) \times 10^{-3}$ s$^{-1}$ to $(2.3 \pm 0.2) \times 10^{-3}$ s$^{-1}$. A paired two-sided $t$-test across biological replicates gave $P = 0.002$, and a permutation test that shuffled treatment labels among recordings within each replicate gave $P < 0.01$ (Supplementary Fig. S4c). NA inhibition therefore reduced the accumulation of local directional bias into arrival more strongly than displacement-level alignment.

### 6 Binder–cleaver heterogeneity and transport functions

The population calculation maps particle architecture and molecular parameters onto exploration, escape from adhesive regions and persistent attachment, showing how a heterogeneous population can span all three.

#### 6.1 Molecular coordinates and function scores

For $N_{\mathrm{tot}} = N_b + N_c$ total ligand sites, we combine binder number and affinity into the average number of attached binders per site, and cleaver number and activity into the cleavage capacity per site:

$$\phi_b = \frac{N_b \nu_b}{N_{\mathrm{tot}}(K_D + \nu_b)}, \qquad \chi_C = \frac{K_C N_c}{N_{\mathrm{tot}}}.$$

Here $\nu_b$ is the number of receptors accessible to one binder at the reference density. Thus $N_{\mathrm{tot}}\phi_b$ is the average number of attached binders. The total cleavage rate is $\lambda_c = k_0\sqrt{\pi}\alpha N_{\mathrm{tot}}\chi_C$. The probability of at least one attachment is $P_{\mathrm{att}} = 1 - [K_D/(K_D + \nu_b)]^{N_b}$, and the probability of cleavage during the architecture-specific persistence time $\tau$ is $P_{\mathrm{cleave}} = 1 - e^{-\lambda_c \tau}$.

For a mixed particle, the uniform theory also gives

$$P_{\mathrm{local}} = \frac{P_0^{\mathrm{dep}} - P_0^{\mathrm{intact}}}{1 - P_0^{\mathrm{intact}}},$$

the probability that local cleavage removes the receptors supporting an existing attachment. Here $P_0^{\mathrm{intact}}$ and $P_0^{\mathrm{dep}}$ are the probabilities that no binder is attached over intact and locally depleted receptors. Spatial separation gives $P_{\mathrm{local}} = 0$ for a polarized particle at this order. Cleavage can therefore break the few contacts of a weakly attached particle or remove the receptors sustaining a more strongly attached mixed particle:

$$P_{\mathrm{esc}} = (1 - P_{\mathrm{att}})P_{\mathrm{cleave}} + P_{\mathrm{att}}P_{\mathrm{local}}.$$

This local escape probability is distinct from the long-time loss of mixed particle mobility already included in $D_{\mathrm{eff}}$.

The positive diffusivity gain and $\mathrm{CI}_{\mathrm{path}}$ define normalized spreading and sensing capacities. Each is multiplied by $1 - P_{\mathrm{esc}}$, and their larger value defines exploration strength. Escape strength is $P_{\mathrm{esc}}$. Persistent attachment defines

$$U_{\mathrm{exploit}} = P_{\mathrm{att}}(1 - P_{\mathrm{local}})\frac{B_t}{\zeta_{t,a}}(1 - P_{\mathrm{cleave}}),$$

scaled once by its maximum over the same parameter range. Here $\zeta_{t,a} = \Gamma_a + B_t$ is the total translational friction and $B_t/\zeta_{t,a}$ is the fraction caused by receptor attachments. The particle friction $\Gamma_a$ is axial for polarized motion and transverse for mixed motion.

#### 6.2 Function maps and population allocation

Main-text Fig. 6a,b shows the largest score available for function $r$ at fixed $(\phi_b, \chi_C)$:

$$U_r^{\max}(\phi_b, \chi_C) = \max_{1 \le N_b \le N_{\mathrm{tot}}-1} U_r(\phi_b, \chi_C, N_b).$$

Different binder allocations can maximize different functions at the same molecular coordinates. In the mixed layout, cleavage breaks the few contacts present at low $\phi_b$, while at high $\phi_b$ local receptor depletion removes otherwise stable contacts. The two escape routes join at strong cleavage.

Main-text Fig. 6c uses a polarized explorer and mixed escape specialist at one shared molecular state, together with a binder-rich polarized exploit specialist. If specialist $r$ has score $U_r$, prescribed demand $d_r$ and population fraction $\pi_r$, we choose the fractions that maximize the minimum coverage $\pi_r U_r / d_r$. For these three specialists,

$$\pi_r^* = \frac{d_r/U_r}{\sum_s d_s/U_s}.$$

Equal demands define the central allocation in main-text Fig. 6c. The displayed sweep varies $d_{\text{explore}}/d_{\text{exploit}}$ from 1/20 to 20 while holding $d_{\text{escape}}$ fixed, moving the population from an exploitation-dominated to an exploration-dominated mixture without removing the escape component. The two endpoints and exact equal-demand solution have explore:escape:exploit fractions 0.045:0.201:0.754, 0.351:0.353:0.296 and 0.789:0.177:0.033, respectively. Each star is projected onto $\lambda_{\text{bc}} = \log_{10}[(\phi_b + 0.003)/(\chi_C + 0.003)]$, where the small offset keeps the logarithm finite at either axis. Explore and escape share the lower mode because they use the same molecular coordinates but different architectures; exploit occupies a binder-rich mode. Each stacked density corresponds to one star in the simplex. The blue, orange and purple areas give the explore, escape and exploit fractions, and the solid outline is their sum. Theory fixes the centers and areas; a common width only displays their overlap.

### References


[1] Vahey, M. D. & Fletcher, D. A. Influenza A virus surface proteins are organized to help penetrate host mucus. *eLife* **8**, e43764 (2019).

[2] Gillespie, D. T. Exact stochastic simulation of coupled chemical reactions. *The Journal of Physical Chemistry* **81**, 2340–2361 (1977).

[3] Ermak, D. L. & McCammon, J. A. Brownian dynamics with hydrodynamic interactions. *The Journal of Chemical Physics* **69**, 1352–1360 (1978).

[4] Fernandes, M. X. & García de la Torre, J. Brownian dynamics simulation of rigid particles of arbitrary shape in external fields. *Biophysical Journal* **83**, 3039–3048 (2002).

[5] Tirado, M. M. & García de la Torre, J. Translational friction coefficients of rigid, symmetric top macromolecules: Application to circular cylinders. *The Journal of Chemical Physics* **71**, 2581–2587 (1979).

[6] Tirado, M. M., López Martínez, C. & García de la Torre, J. Comparison of theories for the translational and rotational diffusion coefficients of rod-like macromolecules: Application to short DNA fragments. *The Journal of Chemical Physics* **81**, 2047–2052 (1984).

[7] Agarwal, S., Veytsman, B., Fletcher, D. A. & Huber, G. Kinetics and optimality of influenza A virus locomotion. *Physical Review Letters* **133**, 248402 (2024).

[8] Schuster, B. S., Suk, J. S., Woodworth, G. F. & Hanes, J. Nanoparticle diffusion in respiratory mucus from humans without lung disease. *Biomaterials* **34**, 3439–3446 (2013).

[9] Parrish, E., Caporizzo, M. A. & Composto, R. J. Network confinement and heterogeneity slows nanoparticle diffusion in polymer gels. *The Journal of Chemical Physics* **146**, 203318 (2017).

[10] Gillespie, D. T. Exact numerical simulation of the Ornstein–Uhlenbeck process and its integral. *Physical Review E* **54**, 2084–2091 (1996).

[11] Morelli, M. J. & ten Wolde, P. R. Reaction brownian dynamics and the effect of spatial fluctuations on the gain of a push–pull network. *The Journal of Chemical Physics* **129**, 054112 (2008).

[12] Pólya, G. Über eine Aufgabe der Wahrscheinlichkeitsrechnung betreffend die Irrfahrt im Straßennetz. *Mathematische Annalen* **84**, 149–160 (1921).

[13] Spitzer, F. Electrostatic capacity, heat flow, and brownian motion. *Zeitschrift für Wahrscheinlichkeitstheorie und Verwandte Gebiete* **3**, 110–121 (1964).

[14] Donsker, M. D. & Varadhan, S. R. S. Asymptotics for the wiener sausage. *Communications on Pure and Applied Mathematics* **28**, 525–565 (1975).
